# Fitness tracking reveals task-specific associations between memory, mental health, and physical activity

**DOI:** 10.1101/2021.10.22.465441

**Authors:** Jeremy R. Manning, Gina M. Notaro, Esme Chen, Paxton C. Fitzpatrick

## Abstract

Physical activity can benefit both physical and mental well-being. Different forms of exercise (e.g., aerobic versus anaerobic; running versus walking, swimming, or yoga; high-intensity interval training versus endurance workouts; etc.) impact physical fitness in different ways. For example, running may substantially impact leg and heart strength but only moderately impact arm strength. We hypothesized that the mental benefits of physical activity might be similarly differentiated. We focused specifically on how different intensities of physical activity might relate to different aspects of memory and mental health. To test our hypothesis, we collected (in aggregate) roughly a century’s worth of fitness data. We then asked participants to fill out surveys asking them to self-report on different aspects of their mental health. We also asked participants to engage in a battery of memory tasks that tested their short and long term episodic, semantic, and spatial memory performance. We found that participants with similar physical activity habits and fitness profiles tended to also exhibit similar mental health and task performance profiles. These effects were task-specific in that different physical activity patterns or fitness characteristics varied with different aspects of memory, on different tasks. Taken together, these findings provide foundational work for designing physical activity interventions that target specific components of cognitive performance and mental health by leveraging low-cost fitness tracking devices.

## Introduction

Engaging in physical activity (exercise) can improve physical fitness by increasing muscle strength [9, 20, 23, 34], bone density [1, 8, 21], cardiovascular performance [24, 32], lung capacity [22, although see [35]], and endurance [43]. Physical activity can also improve mental health [2, 4, 10, 12, 14, 25, 26, 27, 31, 33, 40] and cognitive performance [2, 3, 6, 11].

The physical benefits of exercise can be explained by stress-responses of the affected body tissues. For example, skeletal muscles that are taxed during exercise exhibit stress responses [28] that can in turn affect their growth or atrophy [36]. By comparison, the benefits of physical activity on mental health are less direct. For example, one hypothesis is that physical activity leads to specific physiological changes, such as increased aminergic synaptic transmission and endorphin release, which in turn act on neurotransmitters in the brain [31]. Speculatively, if different physical activity regimens lead to different neurophysiological responses, one might be able to map out a spectrum of signalling and transduction pathways that are impacted by a given type, duration, and intensity of physical activity in each brain region. For example, prior work has shown that physical activity increases acetylcholine levels, starting in the vicinity of the exercised muscles [37]. Acetylcholine is thought to play an important role in memory formation [e.g., by modulating specific synaptic inputs from entorhinal cortex to the hippocampus, albeit in rodents; 30]. Given the central role that these medial temporal lobe structures play in memory, changes in acetylcholine might lead to specific changes in memory formation and retrieval.

In the present study, we hypothesize that (a) different intensities of physical activity will have different, quantifiable impacts on cognitive performance and mental health, and that (b) these impacts will be consistent across individuals. To this end, we collected a year of real-world fitness tracking data from each of 113 participants. We then asked each participant to fill out a brief survey in which they self-evaluated and self-reported several aspects of their mental health. Finally, we ran each participant through a battery of memory tasks, which we used to evaluate their memory performance along several dimensions. We searched the data for potential associations between memory, mental health, and physical activity.

## Methods

We ran an online experiment using the Amazon Mechanical Turk (MTurk) platform [13]. We collected data about each participant’s fitness and physical activity habits, a variety of self-reported measures concerning their mental health, and about their performance on a battery of memory tasks.

### Experiment

#### Participants

We recruited experimental participants by posting our experiment as a Human Intelligence Task (HIT) on the MTurk platform. We limited participation to MTurk Workers who had been assigned a “master worker” designation on the platform, given to workers who score highly across several metrics on a large number of HITs, according to a proprietary algorithm managed by Amazon. One criterion embedded into the algorithm is a requirement that master workers must maintain a HIT acceptance rate of at least 95%. We further limited our participant pool to participants who self-reported that they were fluent in English and regularly used a Fitbit fitness tracker device. A total of 160 workers accepted our HIT in order to participate in our experiment. Of these, we excluded all participants who failed to log into their Fitbit account (giving us access to their anonymized fitness tracking data), encountered technical issues (e.g., by accessing the HIT using an incompatible browser, device, or operating system), or who ended their participation prematurely, before completing the full study. In all, 113 participants contributed usable data to the study.

For their participation, workers received a base payment of $5 per hour (computed in 15 minute increments, rounded up to the nearest 15 minutes), plus an additional performance-based bonus of up to $5. Our recruitment procedure and study protocol were approved by Dartmouth’s Committee for the Protection of Human Subjects. We obtained informed consent using an online form administered to all prospective participants prior to enrolling them in our study. All methods were performed in accordance with the relevant guidelines and regulations.

##### Gender, age, and race

Of the 113 participants who contributed usable data, 77 reported their gender as female, 35 as male, and 1 chose not to report their gender. Participants ranged in age from 19 to 68 years old (25^th^ percentile: 28.25 years; 50^th^ percentile: 32 years; 75^th^ percentile: 38 years). Participants reported their race as White (90 participants), Black or African American (11 participants), Asian (7 participants), Other (4 participants), and American Indian or Alaska Native (3 participants). One participant opted not to report their race.

##### Languages

All participants reported that they were fluent in either 1 or 2 languages (25^th^ percentile: 1; 50^th^ percentile: 1; 75^th^ percentile: 1), and that they were “familiar” with between 1 and 11 languages (25^th^ percentile: 1; 50^th^ percentile: 2; 75^th^ percentile: 3).

##### Reported medical conditions and medications

Participants reported having and/or taking medications pertaining to the following medical conditions: anxiety or depression (4 participants), recent head injury (2 participants), high blood pressure (1 participant), bipolar disorder (1 participant), hypothyroidism (1 participant), and other unspecified conditions or medications (1 participant). Participants reported their current and typical stress levels on a Likert scale as very relaxed (-2), a little relaxed (-1), neutral (0), a little stressed (1), or very stressed (2). The “current” stress level reflected participants’ stress at the time they participated in the experiment. Their responses ranged from -2 to 2 (current stress: 25^th^ percentile: -2; 50^th^ percentile: -1; 75^th^ percentile: 1; typical stress: 25^th^ percentile: 0; 50^th^ percentile: 1; 75^th^ percentile: 1). Participants also reported their current level of alertness on a Likert scale as very sluggish (-2), a little sluggish (-1), neutral (0), a little alert (1), or very alert (2). Their responses ranged from -2 to 2 (25^th^ percentile: 0; 50^th^ percentile: 1; 75^th^ percentile: 2). Nearly all (111 out of 113) participants reported that they had normal color vision, and 15 participants reported uncorrected visual impairments (including dyslexia and uncorrected near- or far-sightedness).

##### Residence and level of education

Participants reported their residence as being located in the suburbs (36 participants), a large city (30 participants), a small city (23 participants), rural (14 participants), or a small town (10 participants). Participants reported their level of education as follows: College graduate (42 participants), Master’s degree (23 participants), Some college (21 participants), High school graduate (9 participants), Associate’s degree (8 participants), Other graduate or professional school (5 participants), Some graduate training (3 participants), or Doctorate (2 participants).

##### Reported water and coffee intake

Participants reported the number of 8 oz cups of water and coffee they had consumed prior to accepting the HIT. Water consumption ranged from 0 to 6 cups (25^th^ percentile: 1; 50^th^ percentile: 3; 75^th^ percentile: 4). Coffee consumption ranged from 0 to 4 cups (25^th^ percentile: 0; 50^th^ percentile: 1; 75^th^ percentile: 2).

#### Tasks

Upon accepting the HIT posted on MTurk, each worker was directed to read and fill out a screening and consent form, and to share access to their anonymized Fitbit data via their Fitbit account. After consenting to participate in our study and successfully sharing their Fitbit data, participants filled out a survey and then engaged in a series of memory tasks (Fig. 1). All stimuli and code for running the full MTurk experiment may be found <u>here</u>.

**Figure 1:**
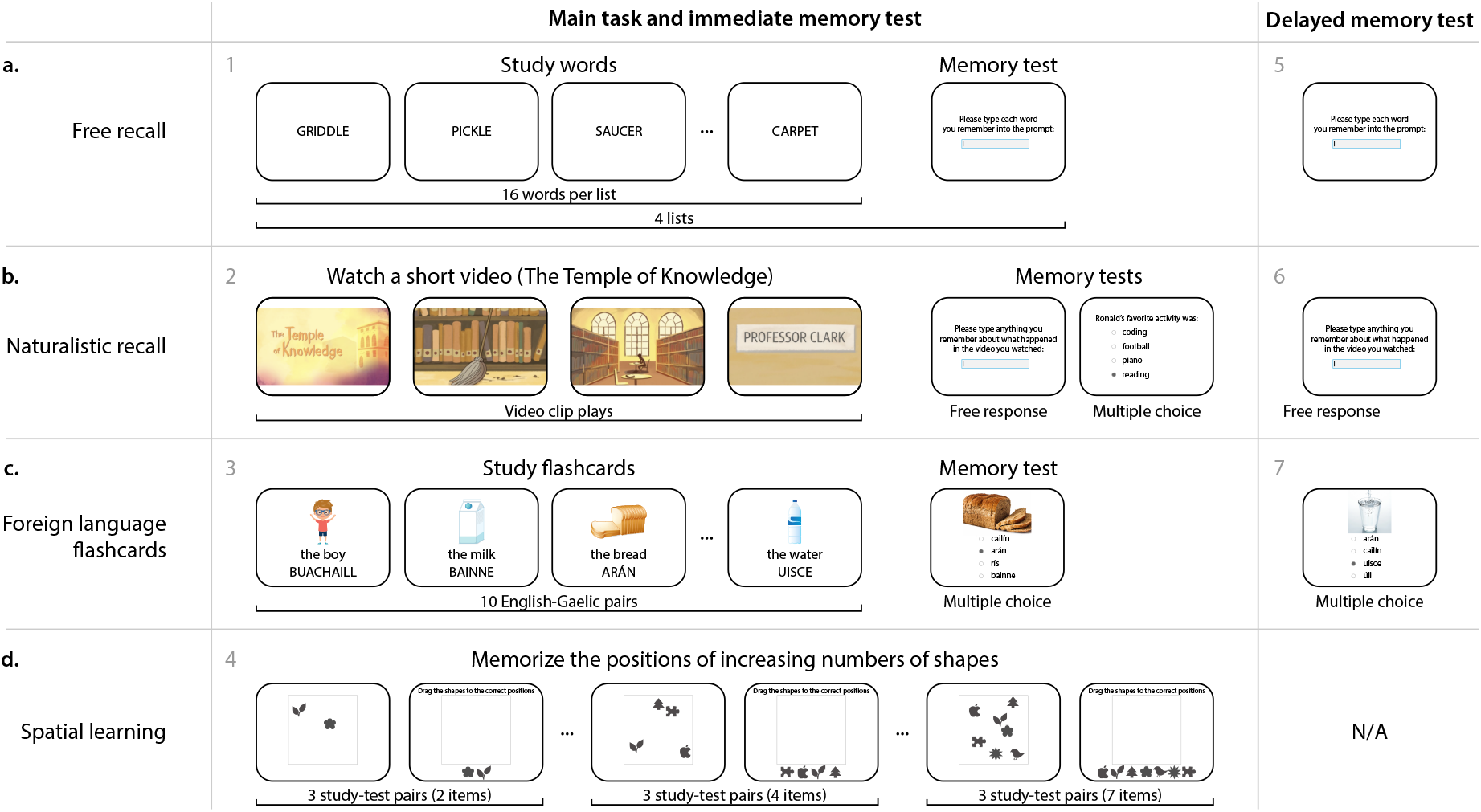
Battery of memory tasks. a. Free recall. Participants study 16 words (presented one at a time), followed by an immediate memory test where they type each word they remember from the just-studied list. In the delayed memory test, participants type any words they remember studying, from any list. **b. Naturalistic recall**. Participants watch a brief video, followed by two immediate memory tests. The first test asks participants to write out what happened in the video. The second test has participants answer a series of multiple choice questions about the conceptual content of the video. In the delayed memory test, participants (again) write out what happened in the video. **c. Foreign language flashcards**. Participants study a sequence of 10 English-Gaelic word pairs, each presented with an illustration of the given word. During an immediate memory test, participants perform a multiple choice test where they select the Gaelic word that corresponds to the given photograph. During the delayed memory test, participants perform a second multiple choice test, where they select the Gaelic word that corresponds to each of a new set of photographs. **d. Spatial learning**. In each trial, participants study a set of randomly positioned shapes. Next, the shapes’ positions are altered, and participants are asked to drag the shapes back to their previous positions. **All panels**. The gray numbers denote the order in which participants experienced each task or test.

##### Survey questions

We collected the following demographic information from each participant: their birth year, gender, highest (academic) degree achieved, race, language fluency, and language familiarity. We also collected information about participants’ health and wellness, including about their vision, alertness, stress, sleep, coffee and water consumption, location of their residence, activity typically required for their job, and physical activity habits.

##### Free recall (Fig. 1a)

Participants studied a sequence of four word lists, each comprising 16 words. After studying each list, participants received an immediate memory test, whereby they were asked to type (one word at a time) any words they remembered from the just-studied list, in any order.

Words were presented for 2 s each, in black text on a white background, followed by a 2 s blank (white) screen. After the final 2 s pause, participants were given 90 s to type in as many words as they could remember, in any order. The memory test was constructed such that the participant could only see the text of the current word they were typing; when they pressed any non-letter key, the current word was submitted and the text box they were typing in was cleared. This was intended to prevent participants from retroactively editing their previous responses.

The word lists participants studied were drawn from the categorized lists reported by [44]. Each participant was assigned four unique randomly chosen lists (in a randomized order), selected from a full set of 16 lists. Each chosen list was then randomly shuffled before presenting the words to the participants. Participants also performed a final delayed memory test where they were given 180 s to type out any words they remembered from *any* of the 4 lists they had studied.

Recalled words within an edit distance of 2 (i.e., a Levenshtein Distance less than or equal to 2) of any word in the wordpool were “autocorrected” to their nearest match. We also manually corrected clear typos or misspellings by hand (e.g., we corrected “hippoptumas” to “hippopotamus”, “zucinni” to “zucchini”, and so on). Finally, we lemmatized each submitted word to match the plurality of the matching wordpool word (e.g., “bongo” was corrected to “bongos”, and so on). After applying these corrections, any submitted words that matched words presented on the just-studied list were tagged as “correct” recalls, and any non-matching words were discarded as “errors.” Because participants were not allowed to edit the text they entered, we chose not to analyze these putative “errors,” since we could not distinguish typos from true misrememberings.

##### Naturalistic recall (Fig. 1b)

Participants watched a 2.5-minute video clip entitled “The Temple of Knowledge.” The video comprises an animated story told to StoryCorps by Ronald Clark, who was interviewed by his daughter, Jamilah Clark. The narrator (Ronald) discusses growing up living in an apartment over the Washington Heights branch of the New York Public Library, where his father worked as a custodian during the 1940s.

After watching the video clip, participants were asked to type out anything they remembered about what happened in the video. They typed their responses into a text box, one sentence at a time. When the participant pressed the return key or typed any final punctuation mark (“.”, “!”, or “?”) the text currently entered into the box was “submitted” and added to their transcript, and the text box was cleared to prevent further editing of any already-submitted text. This was intended to prevent participants from retroactively editing their previous responses. Participants were given up to 10 minutes to enter their responses. After 4 minutes, participants were given the option of ending the response period early, e.g., if they felt they had finished entering all the information they remembered. Each participant’s transcript was constructed from their submitted responses by combining the sentences into a single document and removing extraneous whitespace characters. Following this 4–10-minute free response period, participants were given a series of 10 multiple choice questions about the conceptual content of the story. All participants received the same questions, in the same order. Participants also performed a final delayed memory test, where they carried out the free response recall task a second time, near the end of the testing session. This resulted in a second transcript, for each participant.

##### Foreign language flashcards (Fig. 1c)

Participants studied a series of 10 English-Gaelic word pairs in a randomized order. We selected the Gaelic language both for its relatively small number of native speakers and for its dissimilarity to other commonly spoken languages amongst MTurk workers. We verified (via self report) that all of our participants were fluent in English and that they were neither fluent nor familiar with Gaelic.

Each word’s “flashcard” comprised a cartoon depicting the given word, the English word or phrase in lowercase text (e.g., “the boy”), and the Gaelic word or phrase in uppercase text (e.g., “BUACHAILL”). Each flashcard was displayed for 4 s, followed by a 3 s interval (during which the screen was cleared) prior to the next flashcard presentation.

After studying all 10 flashcards, participants were given a multiple choice memory test where they were shown a series of novel photographs, each depicting one of the 10 words they had learned. They were asked to select which (of 4 unique options) Gaelic word went with the given picture. The 3 incorrect options were selected at random (with replacement across trials), and the orders in which the choices appeared to the participant were also randomized. Each of the 10 words they had learned was tested exactly once.

Participants also performed a final delayed memory test, where they were given a second set of 10 questions (again, one per word they had studied). For this second set of questions participants were prompted with a new set of novel photographs, and new randomly chosen incorrect choices for each question. Each of the 10 original words they had learned were (again) tested exactly once during this final memory test.

##### Spatial learning (Fig. 1d)

Participants performed a series of study-test trials where they memorized the onscreen spatial locations of two or more shapes. During the study phrase of each trial, a set of shapes appeared on the screen for 10 s, followed by 2 s of blank (white) screen. During the test phase of each trial, the same shapes appeared onscreen again, but this time they were vertically aligned and sorted horizontally in a random order. Participants were instructed to drag (using the mouse) each shape to its studied position, and then to click a button to indicate that the placements were complete.

In different study-test trials, participants learned the locations of different numbers of shapes (always drawn from the same pool of 7 unique shapes, where each shape appeared at most one time per trial). They first performed three trials where they learned the locations of 2 shapes; next three trials where they learned the locations of 3 shapes; and so on until their last three trials, where (during each trial) they learned the locations of 7 shapes. All told, each participant performed 18 study-test trials of this spatial learning task (3 trials for each of 2, 3, 4, 5, 6, and 7 shapes).

#### Fitness tracking using Fitbit devices

To gain access to our study, participants provided us with access to all data associated with their Fitbit account from the year (365 calendar days) up to and including the day they accepted the HIT. We filtered out all identifiable information (e.g., participant names, GPS coordinates, etc.) prior to importing their data.

#### Collecting and processing Fitbit data

The fitness tracking data associated with participants’ Fitbit accounts varied in scope and duration according to which device the participant owned (Fig. S1), how often the participant wore (and/or synced) their tracking device, and how long they had owned their device. For example, while all participants’ devices supported basic activity metrics such as daily step counts, only a subset of the devices with heart rate monitoring capabilities provided information about workout intensity, resting heart rate, and other related measures. Across all devices, we collected the following information: heart rate data, sleep tracking data, logged bodyweight measurements, logged nutrition measurements, Fitbit account and device settings, and activity metrics.

##### Heart rate

If available, we extracted all heart rate data collected by participants’ Fitbit device(s) and associated with their Fitbit profile. Depending on the specific device model(s) and settings, this included second-by-second, minute-by-minute, daily summary, weekly summary, and/or monthly summary heart rate information. These summaries include information about participants’ average heart rates, and the amount of time they were estimated to have spent in different “heart rate zones” (rest, out-of-range, fat burn, cardio, or peak, as defined by their Fitbit profile), as well as an estimate of the number of estimated calories burned while in each heart rate zone.

##### Sleep

If available, we extracted all sleep data collected by participants’ Fitbit device(s). Depending on the specific device model(s) and settings, this included nightly estimates of the duration and quality of sleep, as well as the amount of time spent in each sleep stage (awake, REM, light, or deep).

##### Weight

If available, we extracted any weight-related information affiliated with participants’ Fitbit accounts within 1 year prior to enrolling in our study. Depending on their specific device model(s) and settings, this included their weight, body mass index, and/or body fat percentage.

##### Nutrition

If available, we extracted any nutrition-related information affiliated with participants’ Fitbit accounts within 1 year prior to enrolling in our study. Depending on their specific account settings and usage behaviors, this included a log of the specific foods they had eaten (and logged) over the past year, and the amount of water consumed (and logged) each day.

##### Account and device settings

We extracted any settings associated with participants’ Fitbit accounts to determine (a) which device(s) and model(s) are associated with their Fitbit account, (b) time(s) when their device(s) were last synced, and (c) battery level(s).

##### Activity metrics

If available, we extracted any activity-related information affiliated with participants’ Fitbit accounts within 1 year prior to enrolling in our study. Depending on their specific device model(s) and settings, this included: daily step counts; daily amount of time spent in each activity level (sedentary, lightly active, fairly active, or very active, as defined by their account settings and preferences); daily number of floors climbed; daily elevation change; and daily total distance traveled.

##### Comparing recent versus baseline measurements

We were interested in separating out potential associations between *absolute* fitness metrics and *relative* metrics. To this end, in addition to assessing potential raw (absolute) fitness metrics, we also defined a simple measure of recent changes in those metrics, relative to a baseline:

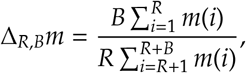

where *m*(*i*) is the value of metric *m* from *i* − 1 days prior to testing (e.g., *m*(1) represents the value of *m* on the day the participant accepted the HIT, and *m*(10) represents the value of *m* 9 days prior to accepting the HIT). We set *R* = 7 and *B* = 30. In other words, to estimate recent changes in any metric *m*, we divided the average value of *m* taken over the prior week by the average value of *m* taken over the 30 days before that.

#### Exploratory correlation analyses

We used a bootstrap procedure to identify reliable correlations between different memory-related, fitness-related, and demographic-related variables. For each of *N* = 10, 000 iterations, we selected (with replacement) a sample of 113 participants to include. This yielded, for each iteration, a sampled “data matrix” with one row per sampled participant and one column for each measured variable. When participants were sampled multiple times in a given iteration, as was often the case, this matrix contained duplicate rows. Next, we computed the Pearson’s correlation between each pair of columns. This yielded, for each pair of columns, a distribution of *N* bootstrapped correlation coefficients. If 97.5% or fewer of the coefficients for a given pair of columns had the same sign, we excluded the pair from further analysis and considered the expected correlation between those columns to be undefined. If > 97.5% of the coefficients for a given pair of columns had the same sign (corresponding to a bootstrap-estimated two-tailed *p* threshold of 0.05), we computed the expected correlation coefficient as:

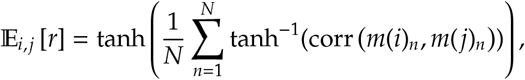

where *m*(*x*)_*n*_ represents column *x* of the bootstrapped data matrix for iteration *n*, tanh is the hyperbolic tangent, and tanh^−1^ is the inverse hyperbolic tangent. We estimated the corresponding *p*-values for these correlations as one minus the proportion of bootstrapped correlations with the same sign, multiplied by two.

#### Reverse correlation analyses

We sought to characterize potential associations between the *dynamics* of participants’ fitnessrelated activities leading up to the time they participated in a memory task and their performance on the given task. For each fitness-related variable, we constructed a timeseries matrix whose rows corresponded to timepoints (sampled once per day) leading up to the day the participant accepted the HIT for our study, and whose columns corresponded to different participants. These matrices often contained missing entries, since different participants’ Fitbit devices tracked fitness-related activities differently. For example, participants whose Fitbit devices lacked heart rate sensors would have missing entries for any heart rate-related variables. Or, if a given participant neglected to wear their fitness tracker on a particular day, the column corresponding to that participant would have missing entries for that day. To create stable estimates, we smoothed the timeseries of each fitness measure using a sliding window of 1 week. In other words, for each fitness measure, we replaced the “observed value” for each day with the average values of that measure (when available) over the 7-day interval ending on the given day.

In addition to this set of matrices storing timeseries data for each fitness-related variable, we also constructed a memory performance matrix, *M*, whose rows corresponded to different memoryrelated variables, and whose columns corresponded to different participants. For example, one row of the memory performance matrix reflected the average proportion of words (across lists) that each participant remembered during the immediate free recall test, and so on.

Given a fitness timeseries matrix, *F*, we computed the weighted average and weighted standard error of the mean of each row of *F*, where the weights were given by a particular memory-related variable (row of *M*). For example, if *F* contained participants’ daily step counts, we could use any row of *M* to compute a weighted average across any participants who contributed step count data on each day. Choosing a row of *M* that corresponded to participants’ performance on the naturalistic recall task would mean that participants who performed better on the naturalistic recall task would contribute more to the weighted average timeseries of daily step counts. Specifically, for each row, *t*, of *F*, we computed the weighted average (across the *S* participants) as:

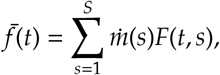

where 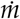 denotes the normalized min-max scaling of *m* (the row of *M* corresponding to the chosen memory-related variable):

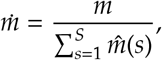

where

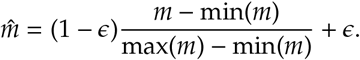

Here, *ϵ* provides a lower bound on the influence of the lowest-weighted participant’s data. We defined *ϵ* = 0.001, ensuring that the lowest-weighted participant had relatively low (but non-zero) influence. We computed the weighted standard error of the mean as:

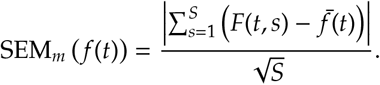

When a given row of *F* was missing data from one or more participants, those participants were excluded from the weighted average for the corresponding timepoint and the weights (across all remaining participants) were re-normalized to sum to 1. The above procedure yielded, for each memory variable, a timeseries of weighted average (and weighted standard error of the mean) fitness tracking values leading up to the day of the experiment.

## Results

Before testing our main hypotheses, we examined the behavioral data from each of four memory tasks (Fig. 1): a random word list learning “free recall” task; a naturalistic recall task whereby participants watched a short video and then recounted the narrative; a foreign language “flashcards” task; and a spatial learning task. Each of the first three tasks (free recall, naturalistic recall, and the flashcards task) included both an immediate (short term) memory test and a delayed (long term) memory test. The spatial learning task included only an immediate test. Participants in all four tasks exhibited general trends and tendencies that have been previously reported in prior work. We were also interested in characterizing the variability in task performance across participants. For example, if all participants exhibited near-identical behaviors or performance on a given task, we would be unable to identify how memory performance on that task varied with mental health or physical activity.

When participants engaged in free recall of random word lists, they displayed strong primacy and recency effects [29] on the immediate memory tests (as reflected by improved memory for early and late list items; Fig. 2a, left and right panels). On the delayed memory test, the recency effect was substantially diminished (Fig. 3a, left and right panels), consistent with myriad previous studies [for review see 18]. Participants also tended to cluster their recalls according to the words’ study positions [17] on both the immediate (Fig. 2a, middle panel) and delayed (Fig. 3a, middle panel) memory tests.

**Figure 2:**
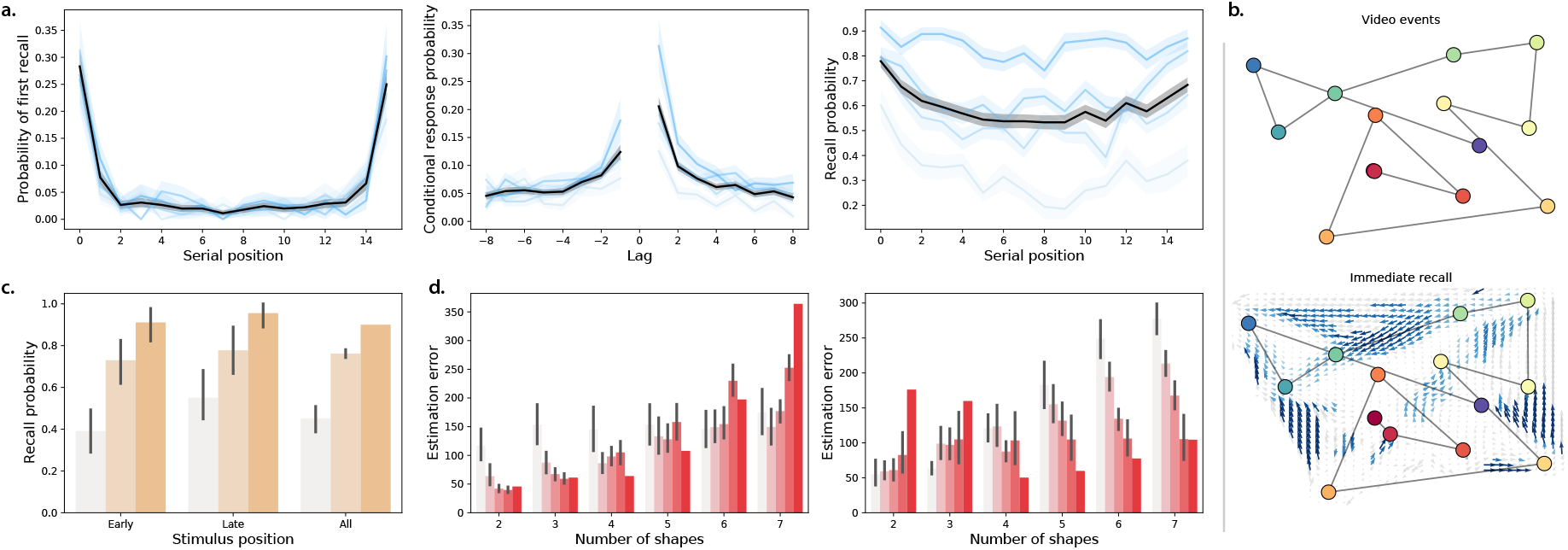
Immediate memory tests. a. Free recall. Left: probability of recalling each word first as a function of its presentation position. Middle: probability of transitioning between successively recalling the word presented at position *i*, followed by word presented at position *i* + Lag. Right: probability of recalling each word as a function of its presentation position. See Figure S2 for additional details. **b. Naturalistic recall**. Top: 2D embedding of a 2.5-min video clip; each dot reflects a narrative event (red denotes early events and blue denotes later events). Bottom: 2D embedding of the averaged transcripts of participants’ recountings of the narrative (dots: same format as top panel). The arrows denote the average trajectory directions through the corresponding region of text embedding space, for any participants whose recountings passed through that region. Blue arrows denote statistically reliable agreement across participants (*p* < 0.05, corrected). See Figure S3 for additional details. **c. Foreign language flashcards**. Each bar denotes the average proportion of correctly recalled Gaelic-English word pairs from early (first 3), late (last 3), or all (i.e., all 10) study positions. See Figure S4 for additional details. **d. Spatial learning**. Average estimation error in shape locations as a function of the number of shapes. See Figure S5 for additional details. All panels: error bars and error ribbons denote bootstrap-estimated 95% confidence intervals. Shading (saturation) denotes results for different subsets of participants assigned based on their task performance (Figs. S2, S3, S4, and S5 provide information about which performance metrics and values the shading reflects; in general more saturated colors denote participants who performed better on the given task.) In Panel d, participants are grouped in two ways; in the left panel, participants are grouped according to the *y*-intercepts of regression lines (estimation error as a function of the number of shapes); in the right panel, participants are grouped according to the slopes of the same regression lines.

**Figure 3:**
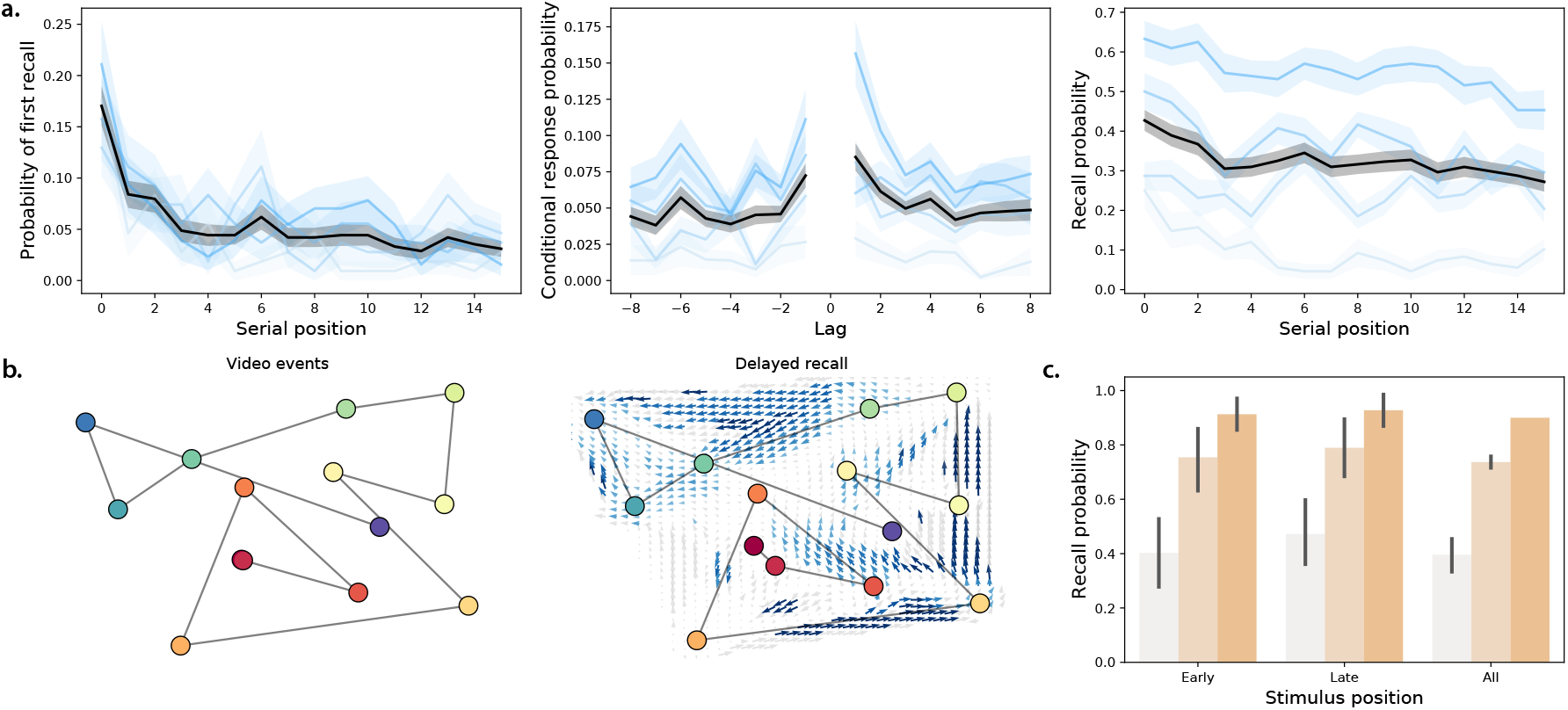
Delayed memory tests. a. Free recall. These panels are in the same format as Figure 2a, but they reflect performance on the delayed free recall task. For additional details see Figure S2. **b. Naturalistic recall**. These panels are in the same format as Figure 2b, but the right panel reflects performance on the delayed naturalistic recall task. For additional details see Figure S3. **c. Foreign language flashcards**. This panel is in the same format as Figure 2c, but it reflects performance on the delayed flashcards test. For additional details see Figure S4.

When participants engaged in naturalistic recall by recounting the narrative of a short story video, they reliably and accurately remembered the major narrative events on both the immediate (Fig. 2b) and delayed (Fig. 3b) tests. This is consistent with prior work showing that memory for rich narratives is both detailed and accurate [7, 15].

Performance on the foreign language flashcards task (immediate: Fig. 2c; delayed: Fig. 3c) varied substantially across participants, and did not show any clear serial position effects. Participants also displayed substantial variation in performance on the spatial learning task (Fig. 2d). In general, participants reported the shape’s positions more accurately when there were fewer shapes. However, both the baseline estimation accuracy and the rate of decrease in accuracy as a function of increasing number of memorized locations varied substantially across participants.

In addition to observing substantial across-participant variability in memory performance, we also observed substantial variability in participants’ fitness and activity metrics (Fig. 4). We examined recent measurements, averaged over the week prior to testing (Fig. 4a), baselined measurements (average over the prior week, divided by the average over the preceding 30 days; Fig. 4b), along with more gradually varying measures that tended to remain relatively static over timescales of weeks to months (Fig. 4c). Figure S6 displays across-participant distributions for a broad selection of these measures, and Figures S7, S8, S9, and S10 show different participants’ fitness metrics, broken down by their performance on different memory tasks.

**Figure 4:**
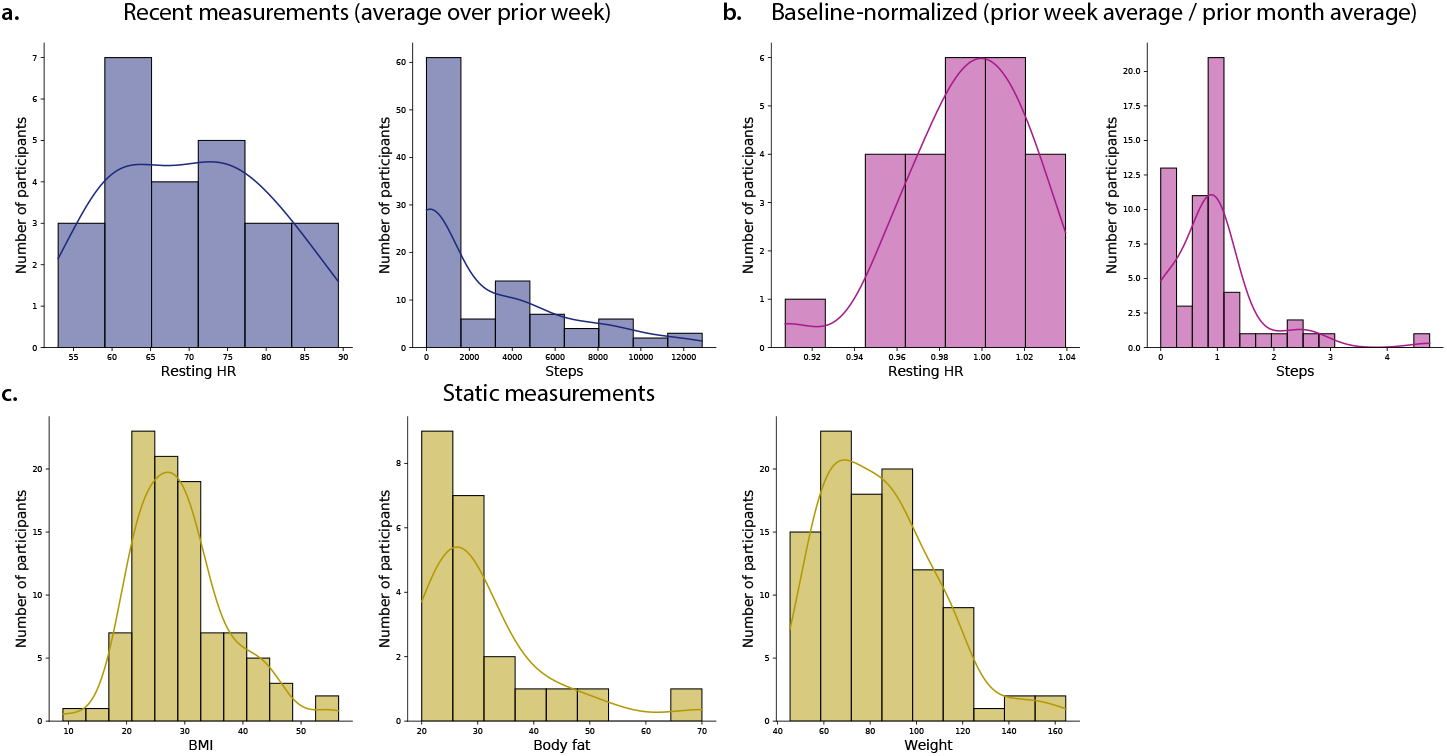
Fitness measures. a. Recent measures. Resting heart rate (HR) and daily step counts, averaged over the week prior to testing. **Baseline-normalized measures**. Resting heart rate and daily step counts averaged over the week prior to testing, divided by the average resting heart rate and step counts averaged over the preceding month. **Static measures**. Body mass index (BMI), body fat percentage, and weight (in kg). For more information see Figures S6, S7, S8, S9, and S10.

We wondered about potential links between the different aspects of participants’ data. For example, if people who engaged in particular intensities of physical activity also tended to perform better on a given memory task, this could suggest that either (a) some property intrinsic to participants who exercised in a particular way might also affect their memory performance on the given task, and/or (b) particular physical activity behaviors could have a causal impact on memory performance. We carried out an exploratory analysis whereby we used a bootstrap-based approach to identify reliable correlations between different aspects of memory performance (Fig. S11), different aspects of fitness (Fig. S12), different demographic attributes (Fig. S13), and correlations between memory performance, fitness information, and demographic attributes (Fig. S14). Specifically, we sought to identify correlations that were present in the same direction (i.e., positive or negative) across different subsets of participants. For each test, we report the average correlation (taken across 10,000 subsets of participants, chosen with replacement) and an associated two-tailed *p*-value, estimated as

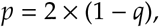

where *q* is the proportion of those 10,000 subsets that exhibited correlations in the same direction (see *Exploratory correlation analyses*). When all 10,000 randomly chosen subsets of participants exhibited correlations in the same direction (i.e., all positive correlations or all negative correlations), we report the *p*-value as *p* < 0.0001.

Several patterns emerged from these analyses. First, we found that participants’ performance on the (within-task) immediate versus delayed memory tests from the free recall, naturalistic recall, and foreign language flashcards tasks were positively correlated (*r*s > 0.25, *p*s < 0.003). This suggests that, within each of these tasks, similar processes or constraints may influence both short term and long term information retrieval. We also found reliable across-task correlations between participants’ (immediate and delayed) performance on the free recall and foreign language flashcards tasks (*r*s > 0.3, *p*s < 0.03).

A large number of fitness-related measures displayed reliable correlations (for a complete report, see Fig. S12). For example, body mass index (BMI) and weight were correlated (*r* = 0.91, *p* < 0.0001). Resting heart rate over the prior week was negatively correlated with recent low-tomoderate-intensity (“fat burn”) cardiovascular activity levels (*r* = 0.70, *p* = 0.0004). Participants’ peak heart rates (averaged over the prior week) were also negatively correlated with recent increases in step counts and daily elevation gains (*r*s < −0.26, *p*s < 0.03), where recent changes were defined as the average values over the seven days leading up to the test day divided by the average values over the preceding 30 days. Several demographic attributes (Fig. S13) displayed trivial correlations (e.g., participants identifying as male never reported identifying as female, and so on). We also observed a negative correlation between reported stress and alertness (*r* = −0.44, *p* < 0.0001), and positive correlations between the reported clarity of the instructions for all tasks (*r*s > 0.26, *p*s < 0.02).

We also found reliable correlations between participants’ fitness and demographic measures and their behaviors in different tasks (Fig. 5; for a complete report, see Fig. S14). For example, recent low-to-moderate-intensity (“fat burn”) cardiovascular activity was positively correlated with immediate (*r* = 0.44, *p* = 0.001) and delayed (*r* = 0.38, *p* = 0.031) recall performance on the naturalistic memory task. Recent sedentary (“out-of-range”) cardiovascular activity was negatively correlated with performance on the spatial learning task (*r* = −0.31, *p* = 0.042), whereas recent high intensity (“peak”) activity was positively correlated with performance on the spatial learning task (*r* = 0.34, *p* = 0.0002). Mental health indicators, such as self-reported stress levels and medications were also associated with differences in memory (Figs. 5a, S14). For example, selfreported stress levels at the time of test were negatively correlated with performance on the delayed memory test for the foreign language flashcards task (*r* = −0.29, *p* = 0.038), whereas participants who were medicated for anxiety and depression tended to perform slightly (but reliably) *better* on the immediate memory test for the foreign language flashcards task (*r* = 0.11, *p* < 0.0001). Mental health indicators were also correlated with several fitness measures (Fig. 5c). For example, participants with higher resting heart rates were less likely to be hypothyroid (*r* = −0.33, *p* < 0.0001). Participants who engaged in more low-intensity (“light”) activity tended to be less anxious and depressed (*r* = −0.12, *p* = 0.03), whereas participants who engaged in more high-intensity activity tended to report higher levels of current (*r* = 0.15, *p* = 0.027) and typical (*r* = 0.21, *p* = 0.012) stress.

**Figure 5:**
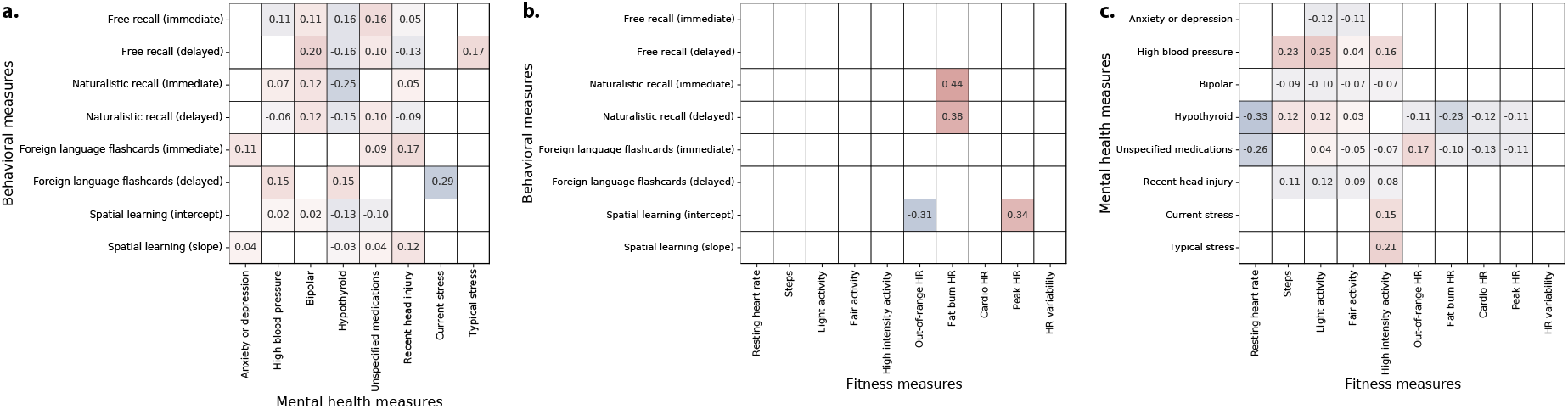
Summaries of correlations between behavioral, fitness, and mental health measures. The reported values in the tables reflect correlations between each pair of measures. Only statistically reliable correlations (*p* < 0.05, corrected) are displayed. **a. Correlations between behavioral and mental health measures**. We adjusted each task’s behavioral measure(s) such that more positive values reflect better performance on the given task. We used participants’ mean recall accuracies to characterize performance on the free recall and foreign language flashcards tasks, and mean precisions to characterize performance on the naturalistic recall tasks. We characterized performance on the spatial learning task using the (inverted and normalized) intercepts and slopes of linear regressions on mean estimation errors as a function of the numbers of studied shapes (also see Figs. 2, 3, S2, S3, S4, and S5). For each mental health measure, more positive values denote greater severity of the given measure. Typical and current stress levels were measured by self report. Mental health information was inferred using each participants’ list of self-reported medications (see *Methods*). Positive correlations indicate that better performance on a given behavioral task is associated with more severe mental health phenotypes. **b. Correlations between fitness and mental health measures**. For each fitness measure, more positive values denote higher observed scores (i.e., higher resting heart rate, more minutes of activity or time spent in each heart rate zone, or greater heart rate variability). The mental health measures in this panel were treated as in Panel a. **c. Correlations between fitness and behavioral measures**. Each measure reflected in this panel was treated as in Panels a and b.

The above analyses indicate that recent differences in fitness-related activity are associated with differences in memory performance and mental health measures. Although the analyses treated these measures on average or in aggregate, many of the measures we collected are dynamic. For example, the amount or intensity of physical activity people engage in can vary over time, and so on. We wondered whether the dynamics of fitness-related measures might relate to memory performance and/or mental health measures. To this end, we carried out a series of reverse correlation analyses (see *Reverse correlation analyses*) to examine whether participants with different cognitive or mental health profiles also tended to display differences in fitness-related measures over time. In particular, we examined fitness data collected from participants’ Fitbit devices over the year prior to their test day in our study. Several example findings are summarized in Figure 6. We found that participants who performed well on the immediate and delayed free recall memory tests and on the naturalistic recall tests tended to be more active than participants who performed poorly on those tests (Figs. 6a, b; S15). Conversely, participants who performed well on the immediate and delayed foreign language flashcards tasks tended to be *less* active. These differences were present even a full year before the testing day. We also found substantial variability across people with different (self-reported) mental health profiles (Figs. 6c, S18). Due to small sample sizes of individuals exhibiting several mental health dimensions, it is difficult to distinguish generalizable trends from individual differences that one or two individuals happened to exhibit. However, several large-sample-size trends emerged. For example, participants who reported higher levels of stress also tended to be slightly more physically active than participants who reported lower stress levels. We found analogous differences in other activity-related measures (Figs. S15 and S18), cardiovascular measures (Figs. S16 and S19), and sleep-related measures (Figs. S17 and S20). Taken together, the analyses suggest that cognitive and mental health differences are also associated with differences in the dynamics of physical health measures.

**Figure 6:**
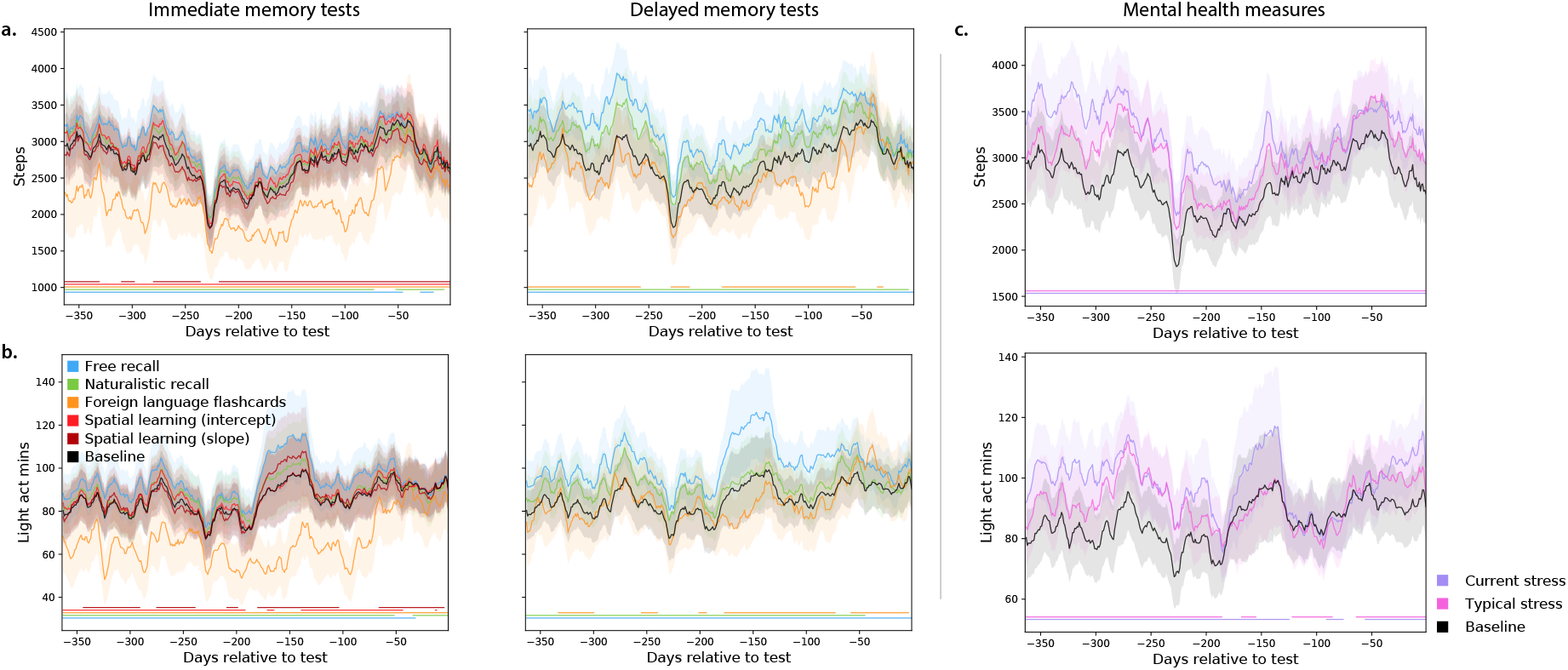
Dynamics of physical activity vary with memory performance and mental health measures. a. Daily step counts. Each timecourse is weighted by either performance on immediate recall tests (left panel) or on delayed recall tests (right panel). The black (baseline) timecourses display the (unweighted) average across all participants. **b. Daily duration (in minutes) of low-intensity physical activity**. Timecourses are displayed in the same format and color scheme as those in Panel A. Analogous timecourses for additional fitness-related measures may be found in Figures S15, S16, and S17. **c. Timecourses of physical activity, weighted by mental health measures**. The timecourses in each panel display the average daily step counts (top panel) or duration of low-intensity activity (bottom panel). The colored lines show average activity dynamics weighted by self-reported stress levels at the start of the experiment (purple) and self-reported “typical” stress levels (pink). The baseline curves (black) display the average across all participants (re-plotted in Panel C to illustrate scale differences across panels). Timecourses for additional mental healthrelated and fitness-related measures may be found in Figures S18, S19, and S20. Error ribbons in all panels denote the standard error of the mean. Horizontal lines below each panel’s timecourses denote intervals over which each weighted measure (color) differs from the unweighted baseline (via a paired sample two-sided *t*-test of the weighted mean values for each measure within a 30-day window around each timepoint; horizontal lines denote *p* < 0.05, corrected).

## Discussion

After collecting a year’s worth of fitness-tracking data from each of 113 participants, we ran each participant in a battery of memory tasks and had them fill out a series of demographic and mental health-related questions. We found that the associations between fitness-related activities, memory performance, and mental health are complex. For example, participants who tended to engage in a particular intensity of physical activity also tended to perform better on some memory tasks but worse on others. This suggests that engaging in one form or intensity of physical activity will not necessarily affect all aspects of cognitive or mental health equally (or in the same direction).

A number of prior studies have shown that engaging in exercise can improve cognitive and mental health [2, 3, 4, 6, 10, 11, 12, 14, 25, 26, 27, 31, 33, 40]. The majority of these studies ask participants in an “exercise intervention” condition (where participants engage in a designated physical activity or training regimen) or a “control” condition (where participants do not engage in the designated activity or training) to perform cognitive tasks or undergo mental health screening. In other words, most primary studies treat “physical activity” as a binary variable that either is or is not present for each participant. Most prior studies also track or manipulate exercise over relatively short durations (typically on the order of days or weeks). Our current work indicates that the true relations between physical activity, cognitive performance, and mental health may be nonmonotonic and heterogeneous across activities, tasks, and mental health measures. These relations can also unfold over much longer timescales than have been previously identified (on the order of months; Fig. 6). However, despite the complexities of the structures of these associations, we also found that they were often remarkably consistent across people. For example, as displayed in Figures 5 and S14, many of the associations between fitness, behavioral, and mental health measures were consistent across over 97.5% of 10,000 randomly chosen subsets of participants.

One important limitation of our study is that we cannot distinguish correlations between different measures from potential causal effects. For example, we cannot know (from our study) whether engaging in particular forms of physical activity *causes* changes in memory performance or mental health, or whether (alternatively) people who tend to engage in similar forms of physical activity also happen to exhibit similar memory and/or mental health profiles. In other words, an overlapping set of processes or person-specific attributes may lead someone to both form particular habits around physical activity and display high or low performance on a given memory test. We do not know whether memory performance or aspects of mental health might be manipulated or influenced by changing the patterns of physical activity someone engages in. For this reason, we have been careful to frame our findings as correlations and associations, rather than to imply knowledge about causal directions of our findings.

Although the present study cannot reveal causal effects, a large prior literature provides some insight into potential causal effects by examining the neural and cognitive effects of a variety of exercise interventions [5, 16, 19, 38, 39, 41, 42]. A limitation of that prior work is that most of these studies examine how relatively short-term changes in physical activity (e.g., on timescales of hours to days or, rarely, weeks to months) affect a cognitive performance on single task or aspect of mental health. The present study examines longer-term physical activity (over a full year), and relates long-term physical activity history to performance on a variety of tasks and to a variety of mental health dimensions.

To the extent that physical activity *does* provide a non-invasive means of manipulating cognitive performance and mental health, our work may have exciting implications for cognitive enhancement. For example, one might imagine building a recommendation system that suggests a particular physical activity regimen tailored to improve a specific aspect of an individual’s cognitive performance (e.g., the efficacy of a student’s study session for an upcoming exam) or mental health (e.g., reducing symptoms of anxiety before an important meeting). Just as strength training may be customized to target a specific muscle group, or to improve performance on a specific physical task, similar principles might also be applied to target specific improvements in cognitive fitness and mental health.

## Acknowledgements

We acknowledge useful discussions with David Bucci, Emily Glasser, Andrew Heusser, Abigail Bartolome, Lorie Loeb, Lucy Owen, and Kirsten Ziman. Our work was supported in part by the Dartmouth Young Minds and Brains initiative, and by NSF grant number 2145172 to J.R.M. The content is solely the responsibility of the authors and does not necessarily represent the official views of our supporting organizations. This paper is dedicated to the memory of David Bucci, who helped to inspire the theoretical foundations of this work. Dave served as a mentor and colleague on the project prior to his passing.

## Data and code availability

All analysis code and data used in the present manuscript may be found here.

## Author contributions

Concept: J.R.M. and G.M.N. Experiment implementation and data collection: G.M.N. Analyses: J.R.M., G.M.N., E.C., and P.C.F. Writing: J.R.M. with input from all authors.

