## Supplemental Materials for "Fitness tracking reveals task-specific associations between memory, mental health, and physical activity"

### *Supplementary materials for:* Fitness tracking reveals task-specific associations between memory, mental health, and exercise

Jeremy R. Manning[1,⋆], Gina M. Notaro[1], Esme Chen[1], and Paxton C. Fitzpatrick[1]

[1]Dartmouth College, Hanover, NH

July 18, 2022

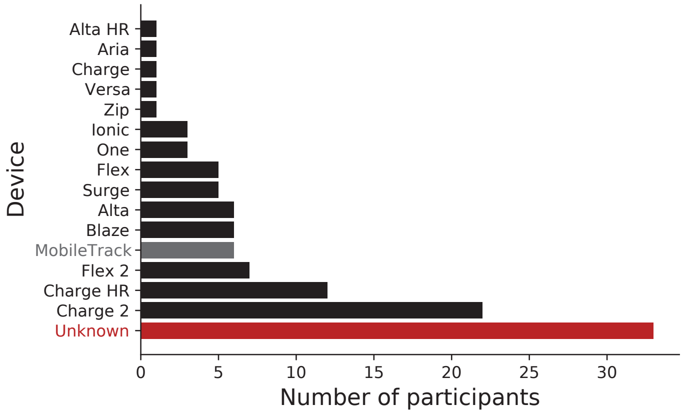


Figure S1: **Fitbit devices.** The bars indicate the numbers of participants whose fitness tracking data came from each model of Fitbit device. “MobileTrack” refers to participants who used smartphone accelerometer information to track their activity via the Fitbit smartphone app. “Unknown” denotes participants whose device information was not specified in their Fitbit data.

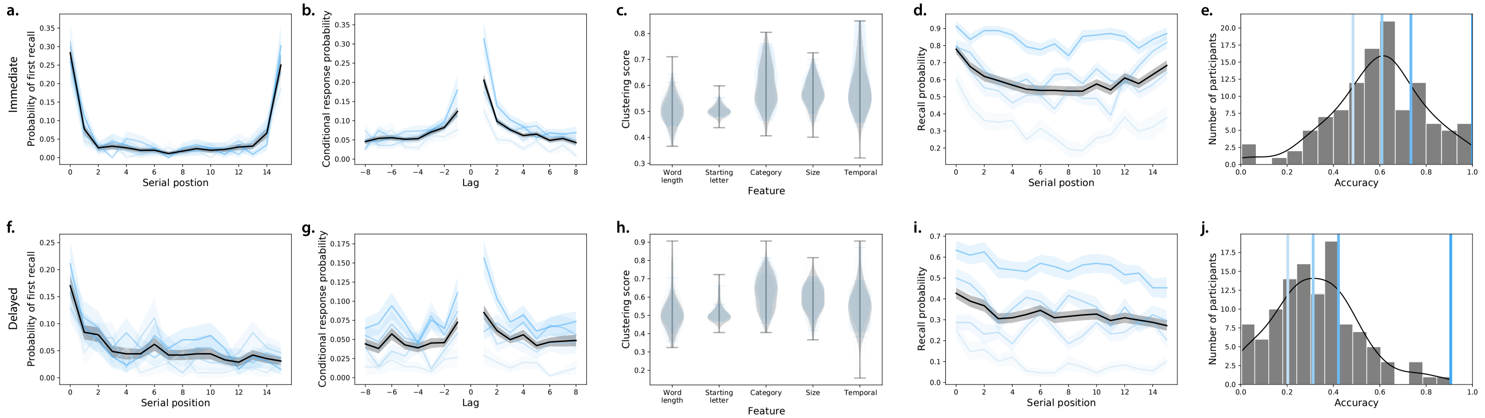


Figure S2: **Free recall behavioral results. a–e. Immediate free recall. a. Probability of first recall.** Probability of recalling each studied word first, as a function of its presentation position. **b. Lag Conditional Response Probability.** Probability of recalling the word presented at position $i$ + Lag following the recall of the word presented at position $i$. **c. Clustering scores.** Each score denotes participants' tendencies to successively recall (cluster) words according to the given feature dimension (Polyn et al., 2009): word length, starting letter, (semantic) category, size (large or small), or presentation position (temporal). **d. Serial position curve.** Probability of recalling each word as a function of its presentation position. **e. Recall accuracy.** Distribution of the average proportions of recalled words, across all lists studied by each participant. **f–j. Delayed free recall.** These panels are in the same formats as Panels a – e, but they reflect performance on the delayed free recall memory tests. All panels: error bars and error ribbons denote bootstrap-estimated 95% confidence intervals. Shading (saturation) denotes results for different subsets of participants, according to the average proportion of words they remembered (group boundaries are indicated by the colored lines in Panels e and j).

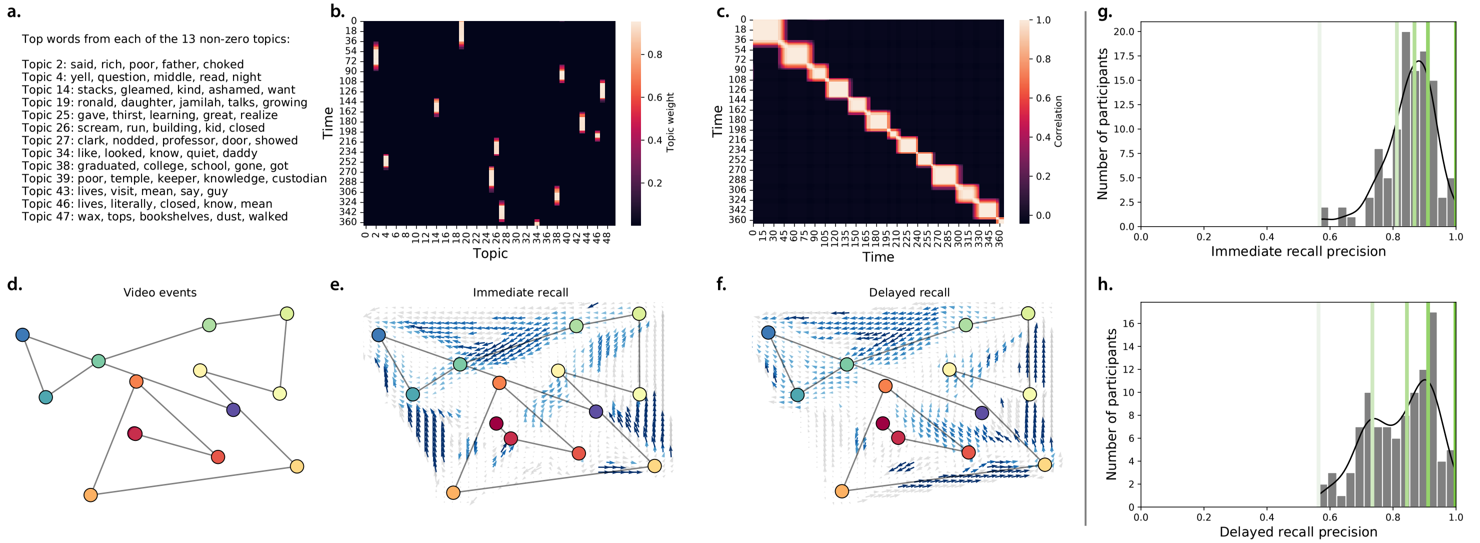


Figure S3: **Naturalistic recall behavioral results. a. Identified topics.** Applying a topic model to a series of sliding windows of the video's transcript (Heusser et al., 2021) revealed a set of 13 unique (non-trivial) topics. The Panel displays the 5 top-weighted words from each topic. **b. Topic timecourse of the video.** Each row displays the topic weights for a single moment of the video. **c. Topic correlation matrix.** The correlations between the topic vectors for each pair of moments from the video reveals an event-like block diagonal structure. **d. Video topic trajectory.** The topic video's topic timecourse (Panel b) has been projected onto 2-dimensions using Uniform Manifold Approximation and Projection (UMAP; McInnes et al., 2018). Each colored dot reflects an event, identified by applying a Hidden Markov Model to the video's topic timecourse (Baldassano et al., 2017; Heusser et al., 2021). Red dots denote earlier timepoints in the video and blue dots denote later timepoints. **e. Immediate recall trajectory.** The black curve displays the average topic timecourse (projected into 2D using UMAP), obtained by applying the topic model shown in Panel A to the participants' written transcripts from the immediate recall test. The arrows denote agreement across participants in the directions of their topic trajectories, for participants whose trajectories intersected the corresponding region of topic space. Blue arrows denote reliable agreement across participants ($p < 0.05$, corrected). **f. Delayed recall trajectory.** This panel is in the same format as Panel e, but displays the trajectory for participants' delayed recall of the video. **g. Immediate recall precision.** Distribution of average recall *precision*, across all the events each participant recalled during the immediate recall test. Precision is defined as the correlation between the topic vector for a given recalled event and the best-matching (most highly correlated) video event's topic vector (Heusser et al., 2021). **h. Delayed recall precision.** This panel is in the same format as Panel g, but displays the average precision values for the delayed memory test.

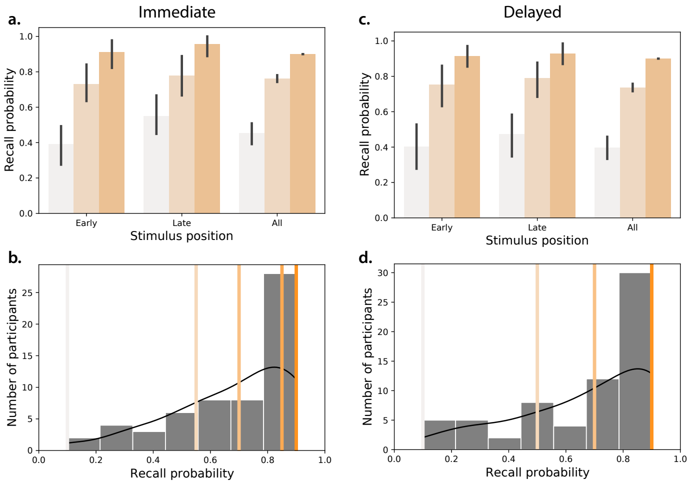


Figure S4: **Foreign language vocabulary learning behavioral results. a. Immediate recall.** Average proportions of correctly identified Gaelic-English word pairs from *early* (first 3) and *late* (last 3) study positions, or aggregated over *all* study positions. **b. Distribution of proportion of correctly recalled pairs on the immediate memory test.** The colored lines indicate quartile boundaries, and correspond to the coloring in Panel a. Note that the right-most edge of the third quartile overlaps with the top quartile, since over 25% of the participants correctly recalled 90% of the word pairs correctly, but no participant achieved 100% accuracy. **c–d. Delayed recall.** These panels are in the same formats as Panels a and b, but reflect performance on the delayed recall test. The error bars in Panels a and c denote bootstrap-estimated 95% confidence intervals.

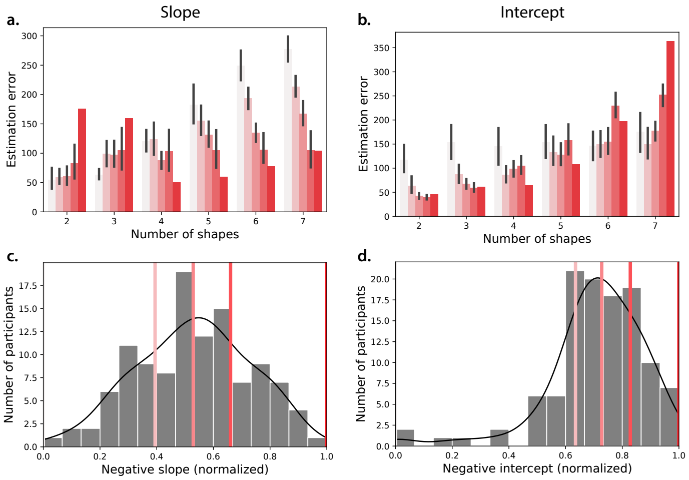


Figure S5: **Spatial learning behavioral results. a–b. Estimation error.** Estimation error is defined as the Euclidean distance between each shape's studied and recalled positions. The bars show the average estimation error (across all shapes for each memory test) versus the number of memorized shapes. **a. Slope.** Estimation errors are broken down by the slopes of regression lines fit to each participant's estimation errors as a function of the number of memorized shapes. Shallower slopes (i.e., smaller decreases in performance associated with memorizing more shapes) are reflected by more saturated coloring. Error bars denote bootstrap-estimated 95% confidence intervals. **b. Intercept.** Bars are in the same format as Panel a, but here estimation errors are broken down by the intercepts of the same regressions used in Panel a. Smaller intercepts (i.e., smaller baseline errors) are reflected by more saturated coloring. **c. Distribution of slopes.** Across-participant distribution of regression slopes. The lines denote quartile boundaries (colors are matched to Panel a). The slopes are multiplied by -1 and normalized to be within the [0, 1] interval such that (adjusted) slopes closer to 1 reflect better performance and slopes closer to 0 reflect worse performance. **b. Distribution of intercepts.** Across-participant distribution of regression intercept terms. The lines denote quartile boundaries (colors are matched to Panel b). The intercepts are multiplied by -1 and normalized to be within the [0, 1] interval such that (adjusted) intercepts closer to 1 reflect better performance and intercepts closer to 0 reflect worse performance.

| Abbreviation | Description | Units |
|---|---|---|
| Resting HR | Average daily resting heart rate | beats per minute (BPM) |
| HR variability | Heart rate variability (average daily variation between heartbeats) | milliseconds (ms) |
| Steps | Number of steps taken each day | steps (count) |
| Distance | Estimated distance traveled each day | kilometers (km) |
| Elevation | Estimated daily elevation gain | feet (ft) |
| Floors climbed | Estimated number of daily floors climbed | floors (count) |
| Light activity | Number of estimated daily minutes spent doing low-intensity activity | minutes (min) |
| Fair activity | Number of estimated daily minutes spent doing moderate-intensity activity | minutes (min) |
| High intensity activity | Number of estimated daily minutes spent doing high-intensity activity | minutes (min) |
| Excess calories | Number of estimated daily calories burned minus estimated calorie intake | calories (cal) |
| Out-of-range HR, OOR mins | Number of daily minutes spent with a heart rate lower than the "fat burn" heart rate zone (50% of the maximum heart rate); maximum heart rate is defined as 220 minus age, in years | minutes (min) |
| Fat burn HR | Number of daily minutes spent with a heart rate between 50% and 69% of the maximum heart rate | minutes (min) |
| Cardio HR | Number of daily minutes spent with a heart rate between 70% and 84% of the maximum heart rate | minutes (min) |
| Peak HR | Number of daily minutes spent with a heart rate between 85% and 100% of the maximum heart rate | minutes (min) |
| Sleep duration | Estimated number of hours slept during the prior 24-hour period | hours (hr) |
| BMI | Body mass index, calculated as the individual's weight (in kg) divided by their height (in m) squared | kilograms per meter squared ($kg/m^2$) |
| Body fat | Estimated percentage of body fat (as a proportion of the individual's total weight) | percentage (%) |
| Weight | Logged or measured weight | kilograms (kg) |
| Water intake | Self-reported water intake | 8 ounce cups (count) |
| Coffee intake | Self-reported coffee intake | 8 ounce cups (count) |

Table S1: **Abbreviations of fitness-related measures.** The abbreviations and descriptions provided in the table apply to the main text and supplement.

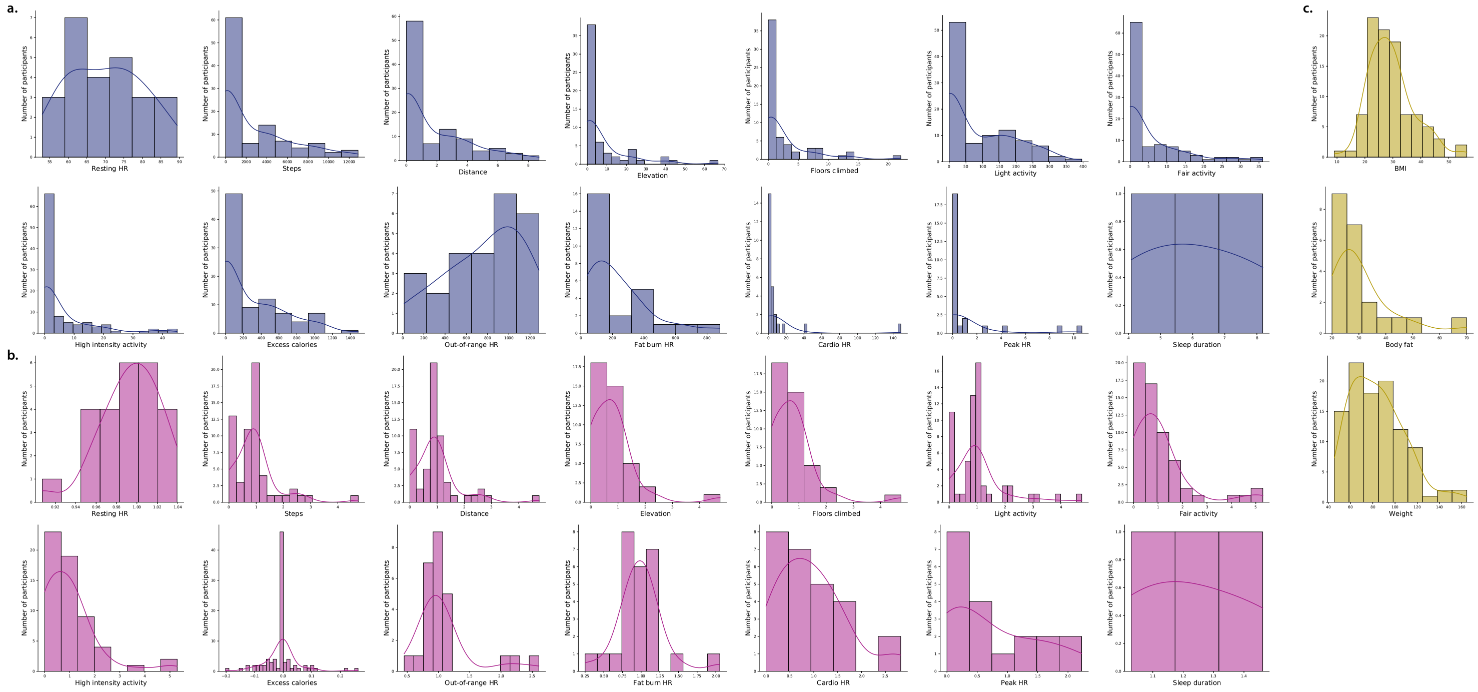


Figure S6: **Distributions of fitness measures.** All abbreviations and measures are described in Table S1. **a. Recent measures.** Daily values of each measure, averaged over the week prior to each participant's test day. **b. Baselined measures.** Daily values of each measure, averaged over the week prior to each participant's test day and divided by the average taken over the preceding 30 days. **c. Static measures.** For each measure, only the most recently logged values are considered. All panels: note that not all measures were captured for all participants. Only participants whose Fitbit records logged each given measure are included in the distribution(s) for that measure.

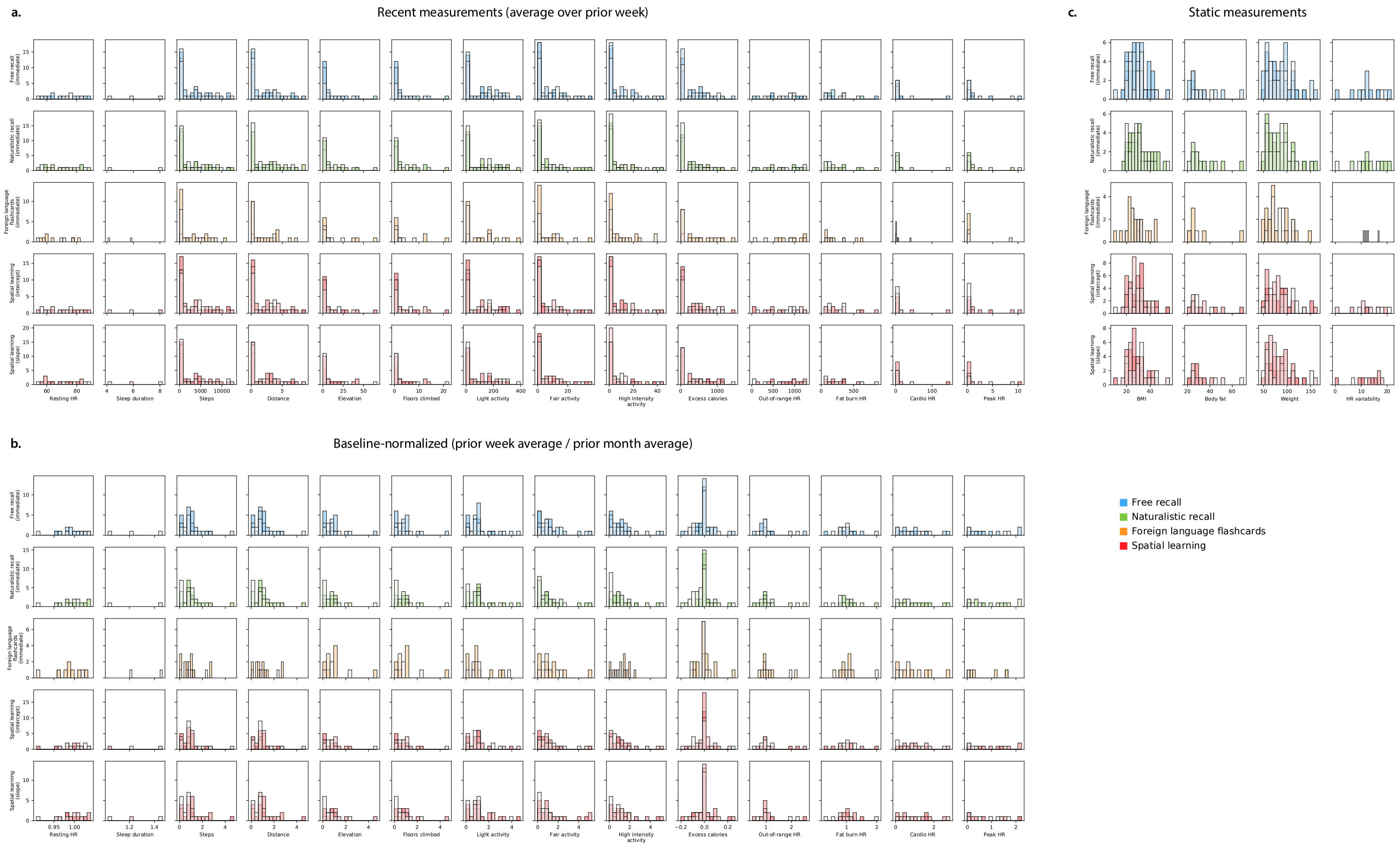


Figure S7: **Distributions of fitness measures, broken down by immediate task performance.** The distributions of recent (Panel a), baselined (Panel b), and static (Panel c) fitness measures (Tab. S1) are broken down by participants' performance on each immediate recall test (Figs. 2, S2, S3, S4, and S5)

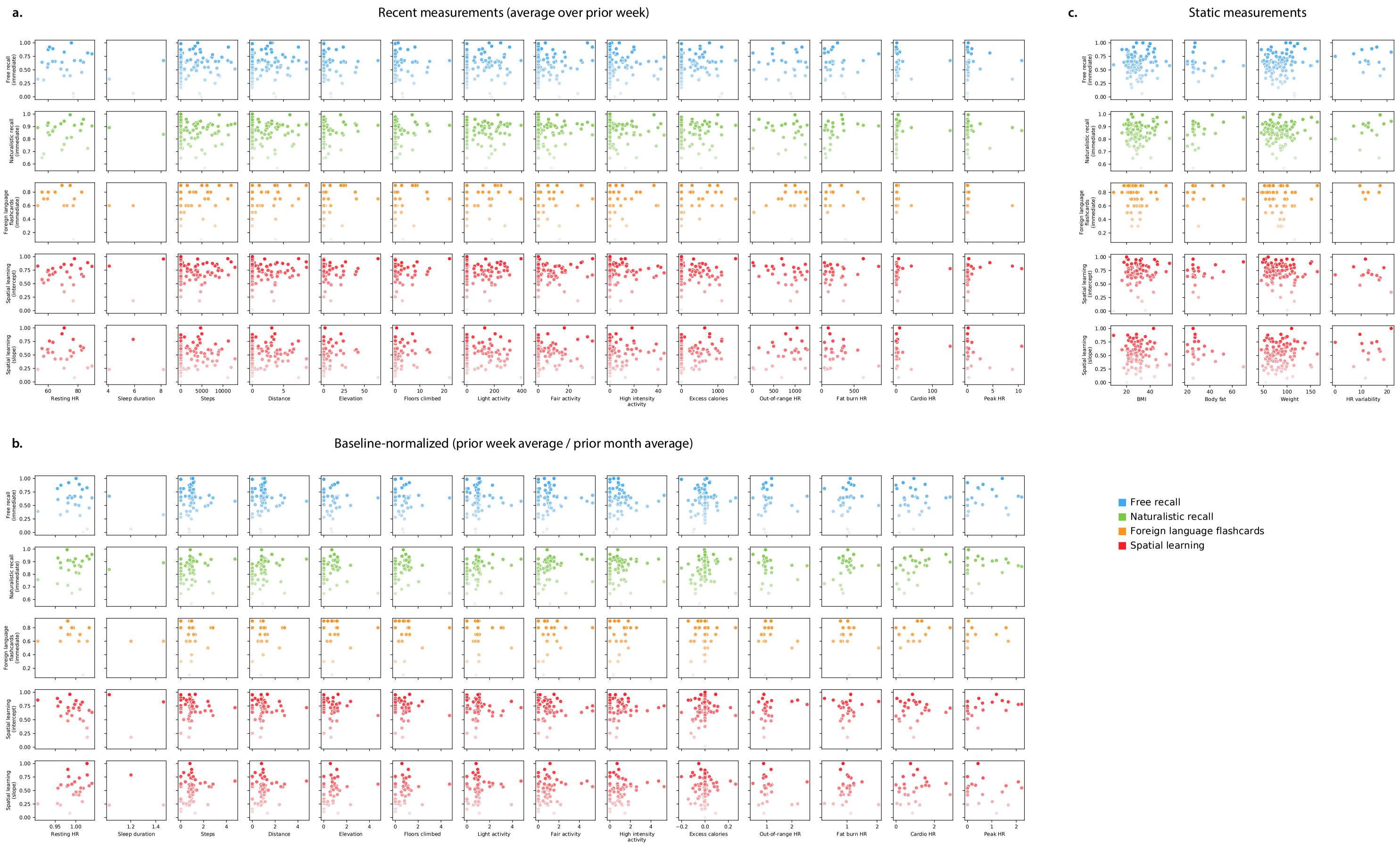


Figure S8: **Scatter plots of fitness measures versus immediate task performance measures.** The scatter plots of recent (Panel a), baselined (Panel b), and static (Panel c) fitness measures (Tab. S1) are plotted against participants' performance on each immediate recall test (Figs. 2, S2, S3, S4, and S5)

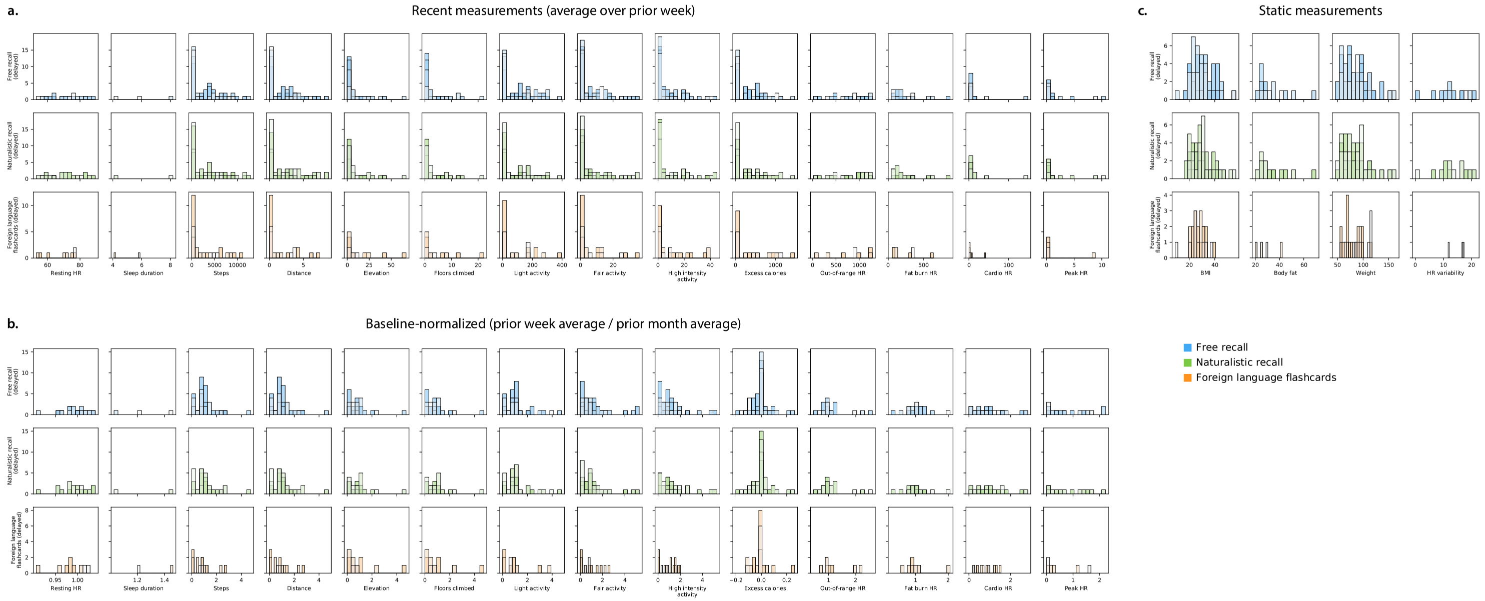


Figure S9: **Distributions of fitness measures, broken down by delayed task performance.** The distributions of recent (Panel a), baselined (Panel b), and static (Panel c) fitness measures (Tab. S1) are broken down by participants' performance on each delayed recall test (Figs. 3, S2, S3, and S4)

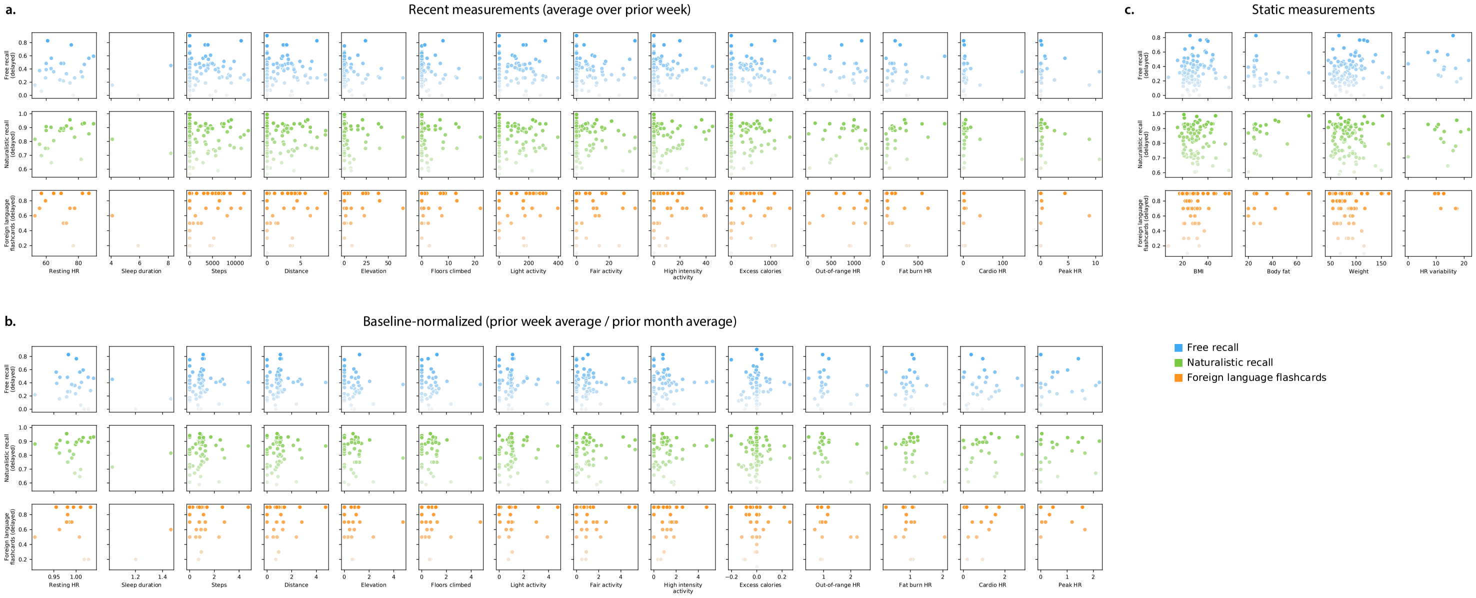


Figure S10: **Scatter plots of fitness measures versus delayed task performance measures.** The scatter plots of recent (Panel a), baselined (Panel b), and static (Panel c) fitness measures (Tab. S1) are plotted against participants' performance on each delayed recall test (Figs. 3, S2, S3, and S4)

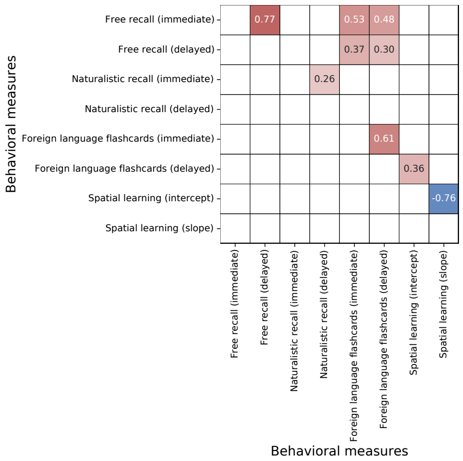


Figure S11: **Bootstrap-estimated reliable correlations between behavioral measures.** The measures are described and summarized in the distributions shown in Figures S2, S3, S4, and S5. The reported values denote the correlation coefficient between the given row and column. Only statistically reliable correlations are displayed ($p < 0.05$, corrected; see *Exploratory correlation analysis*, main text).

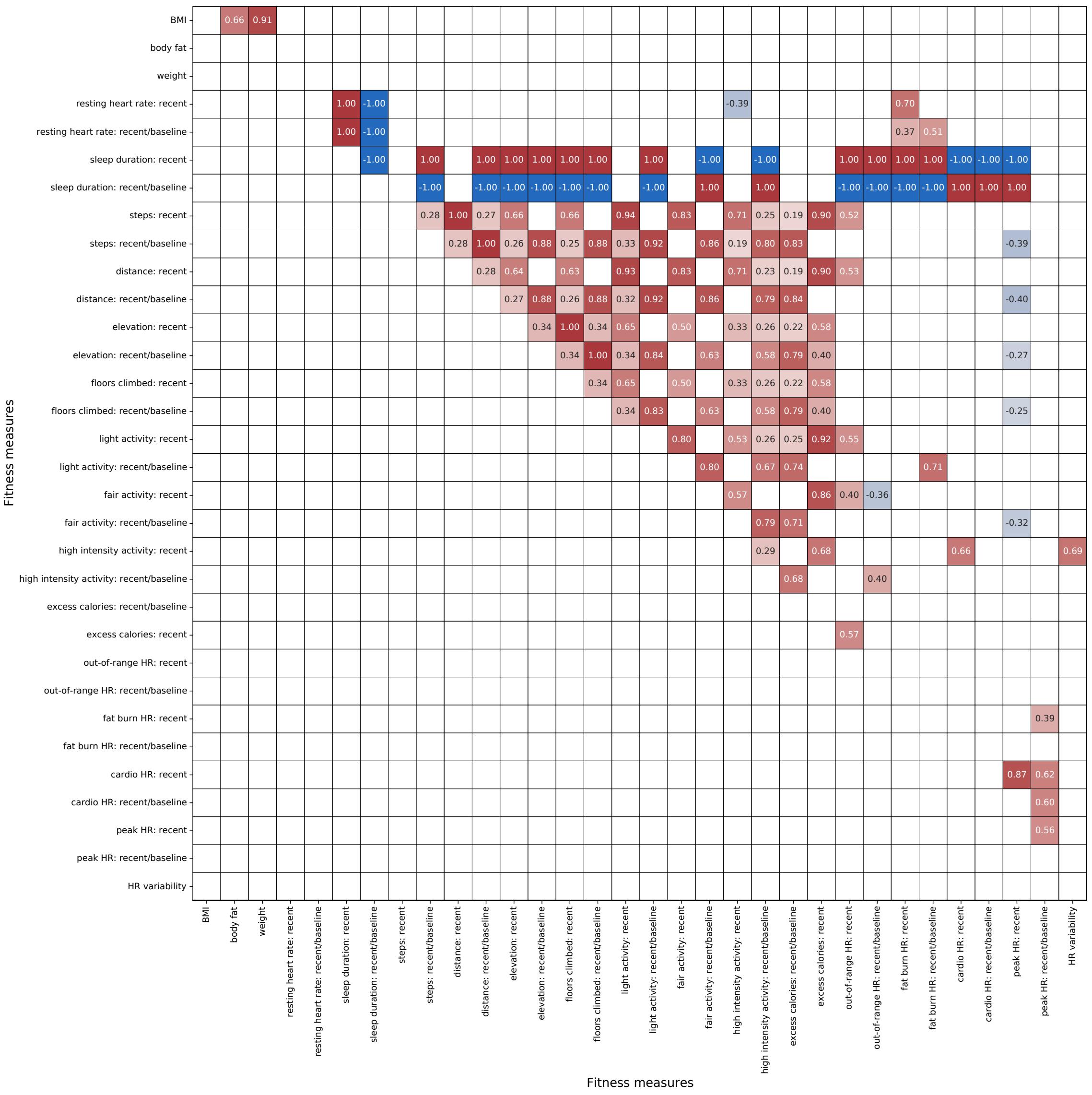


Figure S12: **Bootstrap-estimated reliable correlations between fitness measures.** Fitness measures are described in summarized in Table S1. The reported values denote the correlation coefficient between the given row and column. Only statistically reliable correlations are displayed ($p < 0.05$, corrected; see *Exploratory correlation analysis*, main text).

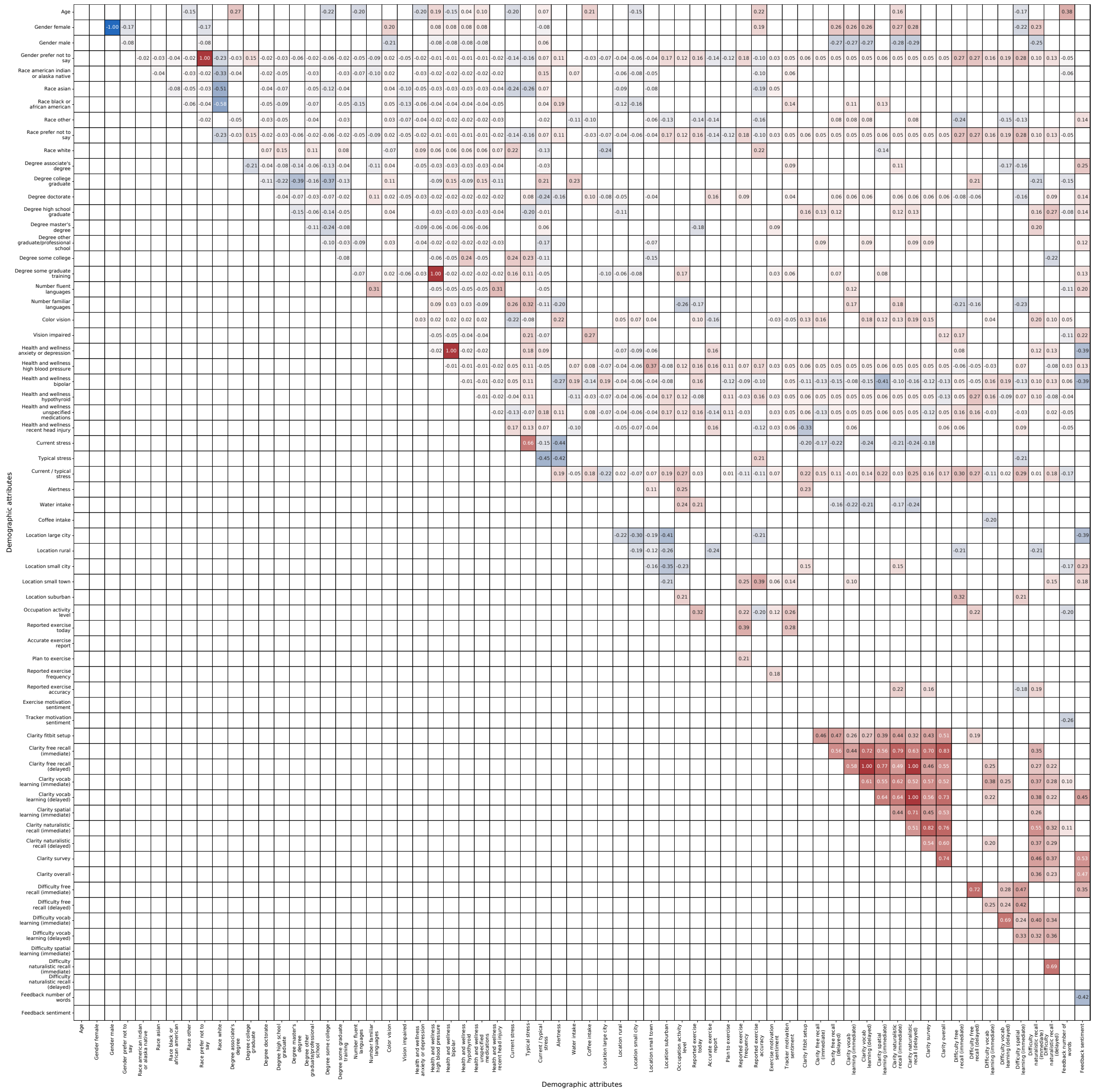


Figure S13: **Bootstrap-estimated reliable correlations between demographic measures.** Demographic measures are described in the main text (*Participants*). The reported values denote the correlation coefficient between the given row and column. Only statistically reliable correlations are displayed ($p < 0.05$, corrected; see *Exploratory correlation analysis*, main text).

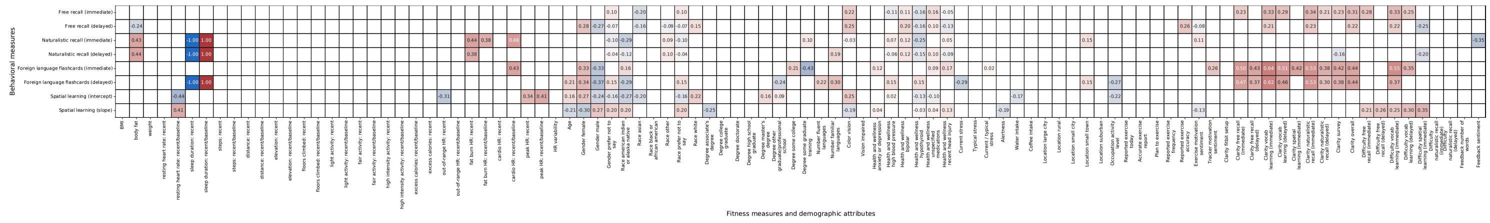


Figure S14: **Bootstrap-estimated reliable correlations between behavioral measures and fitness or demographic measures.** All measures and abbreviations in this figure use the same notations as Figures S11, S12, and S13. Only statistically reliable correlations are displayed ($p < 0.05$, corrected; see *Exploratory correlation analysis*, main text).

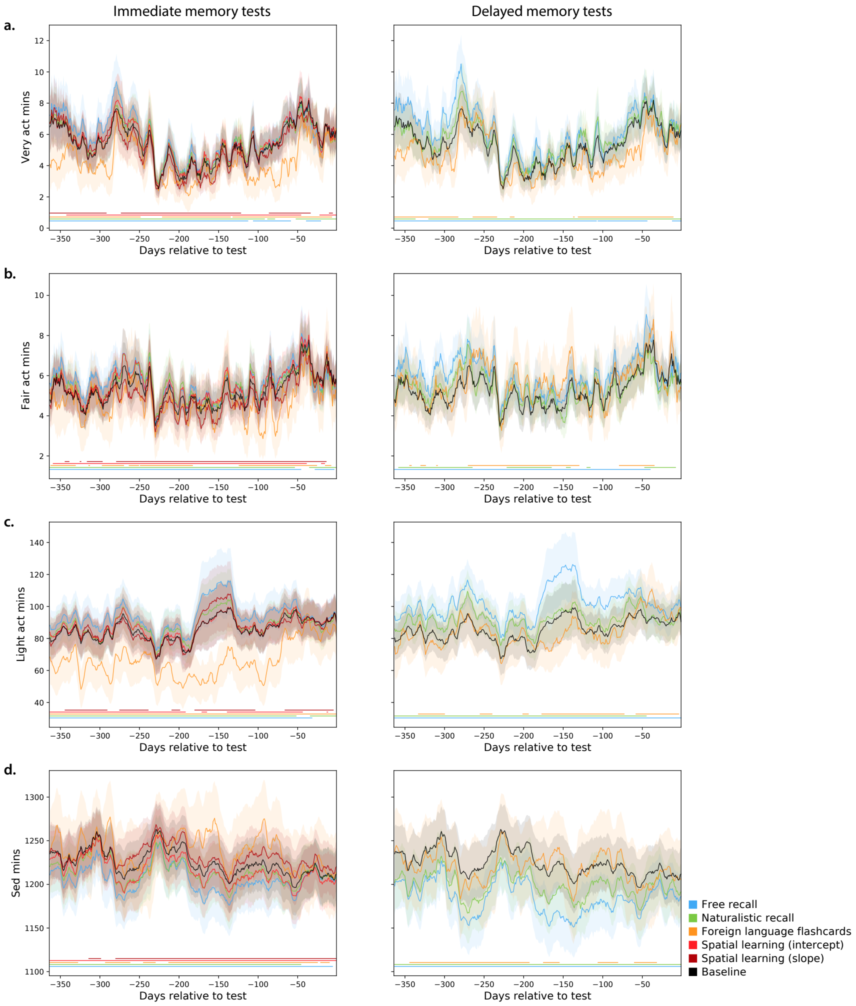


Figure S15: **History of fitness activity levels weighted by behavioral performance.** This figure is in the same format as Figure 6a and b, but shows additional fitness-related measures (Tab. S1).

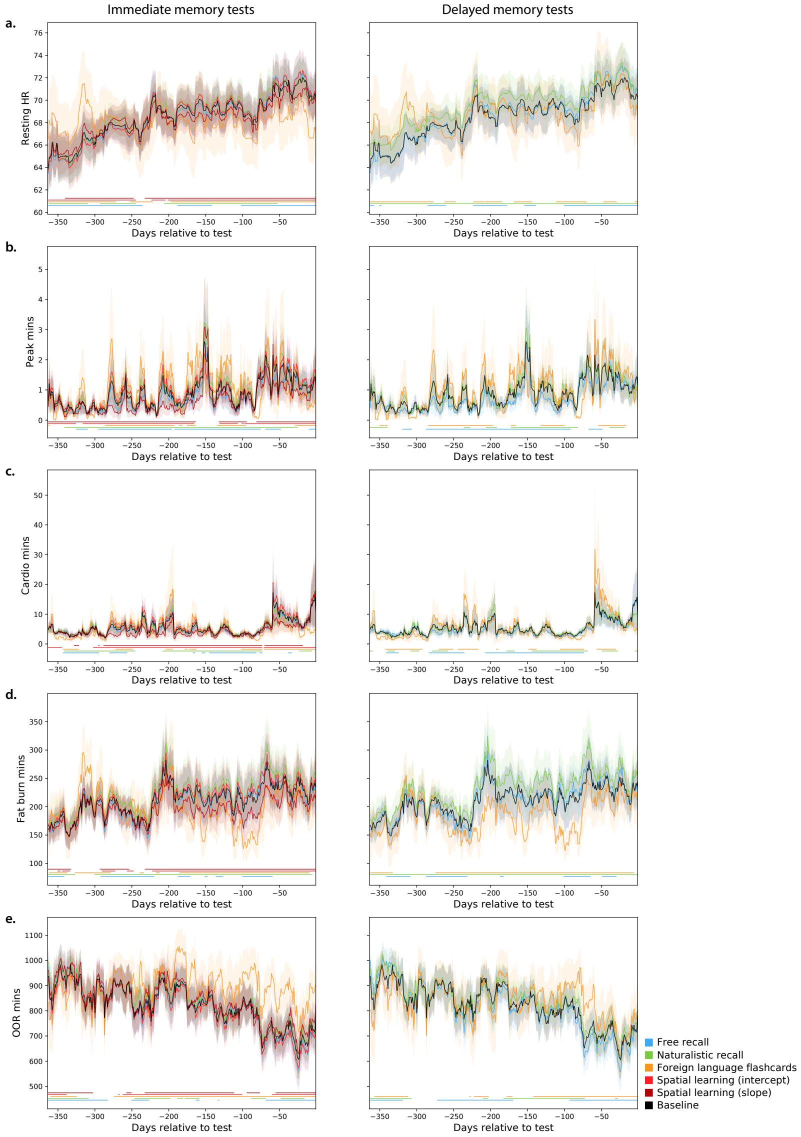


Figure S16: **History of cardiovascular activity weighted by behavioral performance.** This figure is in the same format as Figure 6a and b, but shows additional fitness-related measures (Tab. S1).

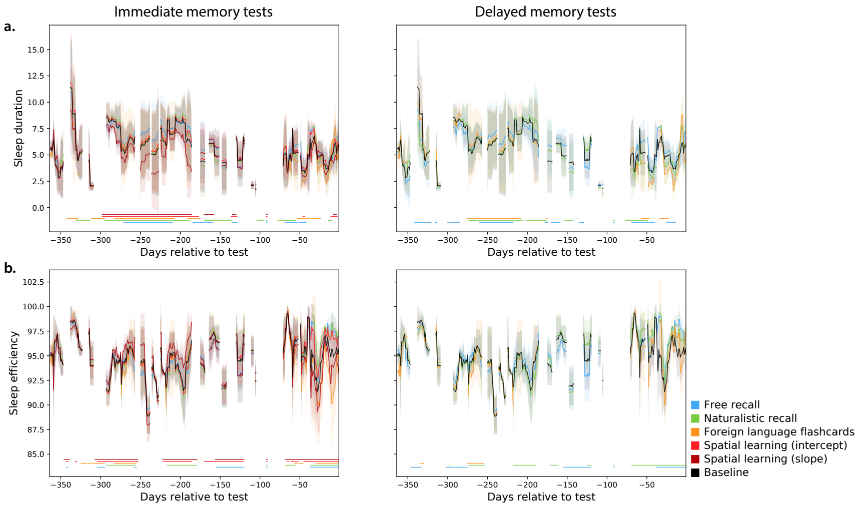


Figure S17: **History of sleep efficiency and duration weighted by behavioral performance.** This figure is in the same format as Figure 6a and b, but shows additional fitness-related measures (Tab. S1).

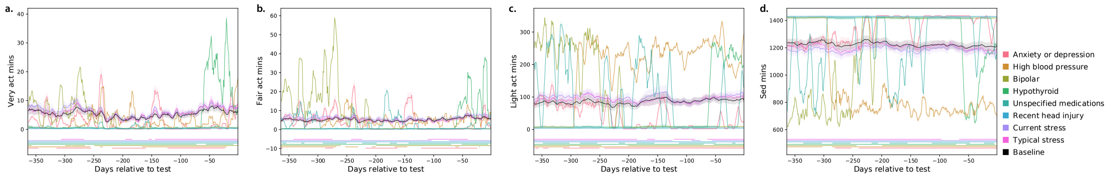


Figure S18: **History of fitness activity levels weighted by mental health factors.** This figure is in the same format as Figure 6c, but shows additional fitness-related and mental health-related measures (Tab. S1).

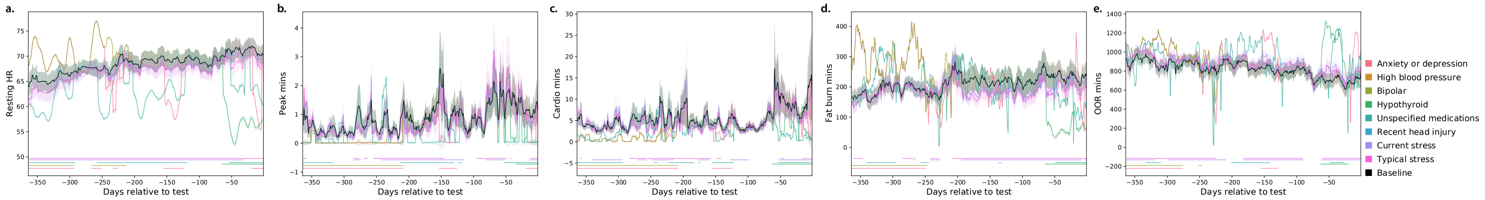


Figure S19: **History of cardiovascular activity weighted by mental health factors.** This figure is in the same format as Figure 6c, but shows additional fitness-related and mental health-related measures (Tab. S1).

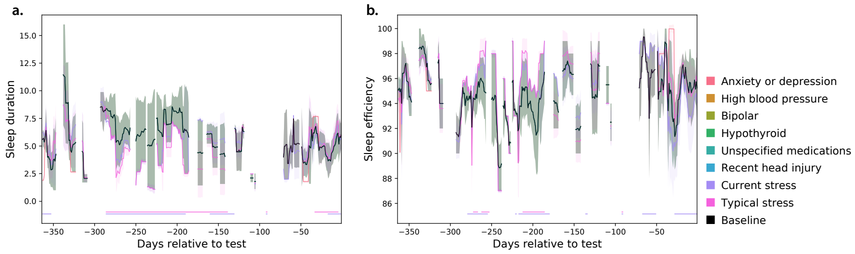


Figure S20: **History of sleep efficiency and duration weighted by mental health factors.** This figure is in the same format as Figure 6c, but shows additional fitness-related and mental health-related measures (Tab. S1).
